# TRBP regulates poly(I:C)-mediated apoptotic pathway by modulating miRNA biogenesis

**DOI:** 10.1101/2024.05.14.594097

**Authors:** Shota Azuma, Yuko Nakano, Tomoko Takahashi, Koji Onomoto, Takeshi Haraguchi, Yutaka Suzuki, Hideo Iba, Mitsutoshi Yoneyama, Yoshimasa Asano, Kumiko Ui-Tei

**Affiliations:** Department of Biological Sciences, Graduate School of Science, The University of Tokyo, Tokyo 113-0033, Japan; Department of Biochemistry and Molecular Biology, Graduate School of Science and Engineering, Saitama University, Saitama 338-8570, Japan; Division of Molecular Immunology, Medical Mycology Research Center, Chiba University, Chiba 260-8673, Japan; Division of RNA Therapy, Medical Mycology Research Center, Chiba University, Chiba, 260-8673, Japan; Department of Computational Biology and Medical Sciences, Graduate School of Frontier Sciences, The University of Tokyo, Chiba 277-8561, Japan

**Keywords:** RNA silencing, TRBP, microRNA, gene network, Virus sensor, LGP2

## Abstract

Interferon (IFN) response is usually triggered by the invasion of viral RNAs, and is known to suppress RNA silencing triggered by microRNAs (miRNAs). However, the detailed mechanism regulating RNA silencing during IFN response has not been elucidated. We previously reported that a virus sensor protein, LGP2, which is an upregulated protein during IFN response, interacts with an RNA-binding protein, TRBP. Since TRBP is an RNA-binding protein which enhances miRNA biogenesis in collaboration with Dicer, the interaction of TRBP with LGP2 inhibits the interaction with Dicer and suppresses the TRBP-mediated miRNA biogenesis. During Sendai virus infection, we found that the inhibition of miRNA maturation caused by TRBP depletion resulted in the induction of apoptosis. However, the global gene expression network regulated by miRNAs during viral infection is not understood. In this study, we analyzed comprehensive data from RNA-sequencing and small RNA-sequencing during the antiviral response induced by poly(I:C) transfection, and investigated how TRBP regulates expression profiles of miRNAs and the downstream mRNAs. A group of miRNAs, including miRNAs preferably bind to TRBP, were downregulated by poly(I:C) transfection. Although about 60 % of miRNAs were situated in the introns of their host genes, the expression levels of mature miRNAs were not correlated with those of host genes. Therefore, the downregulation of the expression levels of mature miRNAs was considered to be not caused by transcriptional inhibition of host genes but by the inhibition of TRBP-mediated miRNA processing. Target prediction analysis suggested that the downregulated miRNAs in the presence of TRBP but not in the absence of TRBP mainly targeted transcription factors. Furthermore, these transcription factors were included in the regulators of apoptosis- and cell cycle-related factors. Some of these regulators were also predicted to be the direct targets of downregulated miRNAs in the presence of TRBP. Thus, it was revealed that the downregulation of miRNAs in the presence of TRBP regulates the expression of apoptosis- and cell cycle-related genes either directly or indirectly through adjusting the expression levels of transcription factors. These results suggest that TRBP has a pivotal role in the determination of cell fates during viral infection regulating the miRNA-mRNA regulatory networks through modifying miRNA biogenesis.

## INTRODUCTION

In mammals, a small non-coding RNA, microRNA (miRNA), plays a critical role in the post-transcriptional regulation of gene expression, fine-tuning various biological processes. MiRNAs are encoded from the genome and become matured form through a series of processing steps. Primary transcripts of miRNAs (pri-miRNAs) are transcribed from the genome by RNA polymerase II (*1*) and processed into precursor miRNAs (pre-miRNAs) by the microprocessor complex, consisting of the nuclear RNase III, Drosha, and a cofactor, DiGeorge syndrome critical region gene 8 (DGCR8) (*2*). The processed pre-miRNAs are exported from the nucleus to the cytoplasm by a transport complex, comprising exportin-5 (XPO5), GTP-binding nuclear protein RAN, and GTP (*3*, *4*). In the cytoplasm, the pre-miRNAs are further processed into mature miRNA duplexes approximately 22 nucleotides (nt) in length by an RNase III enzyme, Dicer (*5*). This process is facilitated by TAR RNA-binding protein (TRBP), interacting with Dicer (*6*). The mature miRNA duplex is loaded onto Argonaute (AGO) protein in the RNA-induced silencing complex (RISC)-loading complex and unwound into single-stranded RNAs (*6*, *7*). One of these two RNA strands, termed as a miRNA guide strand, is remained on AGO protein to form an active RISC and functions as a guide to suppress target mRNA (*8*, *9*). The miRNA guide strand recognizes target mRNAs by partial complementarities in nucleotide sequences. Then, the expressions of target genes are suppressed by cleavage or translational repression of their mRNAs.

Recently, we discovered that the post-transcriptional regulation by miRNAs plays critical roles in antiviral response through regulating apoptosis (*10*). During virus infection, miRNAs are known to regulate both virus-derived RNAs and endogenous RNAs, which leads to regulating various antiviral responses. In mammals, the primary defense system against viral infection is the innate immune response which includes interferon (IFN) response. During the IFN response, antiviral genes such as protein kinase R (PKR), 2’-5’-oligoadenylate synthetase (OAS), or retinoic acid-inducible gene-I (RIG-I) like receptors (RLRs) are known to be upregulated (*11*). RLRs are cytoplasmic virus sensor proteins, which include RIG-I (also known as DDX58), melanoma differentiation-associated gene 5 (MDA5) (also known as interferon induced with helicase C domain 1, IFIH1), and laboratory of genetics and physiology 2 (LGP2) (also known as DHX58) (*12*). RIG-I and MDA5 play a significant role in producing type-I IFN by transmitting signals to IFN-β promoter stimulator 1 (IPS-1) (also known as mitochondrial antiviral signaling protein, MAVS) via its caspase recruitment domain (CARD) (*13*). LGP2 lacks the CARD domain, but binds to double-stranded RNA (dsRNA) much more strongly than RIG-I and MDA5 (*13*). Therefore, LGP2 is not considered to regulate IFN signaling like other virus sensor proteins, and its detailed function has not been fully elucidated. We recently discovered that LGP2 interacts with an RNA silencing enhancer, TRBP (*10*, *14*). TRBP is a dsRNA binding protein (dsRBP) and plays significant roles in miRNA biogenesis. In RNA silencing, TRBP enhances the processing of a specific type of pre-miRNAs into mature miRNAs (*14*, *15*), by forming a heterodimer with Dicer protein. During IFN response, the formation of the TRBP-Dicer complex is inhibited, because upregulated LGP2 competitively binds to TRBP through its dsRNA binding domains (dsRBDs). The inhibition of maturation of specific types of miRNAs leads to the upregulation of their target mRNAs. During viral infection, it was also reported that TRBP interacts with the upregulated PKR through its dsRBDs and inhibits PKR-mediated translational repression (*16*). Thus, TRBP plays multiple roles during viral infection through its interaction with pre-miRNAs as well as other proteins. However, the detailed impact of TRBP during viral infection is not studied, especially in terms of the global regulation of gene expression profiles of mature miRNAs and their downstream genes.

In this study, we analyzed the miRNA-mRNA regulatory networks during antiviral response. Our previous reports revealed that antiviral responses are induced in wild-type (WT) and TRBP^-/-^ human HeLa cells through Sendai virus (SeV) infection (*10*). To investigate the gene expression network during viral infection, we used poly(I:C) to induce antiviral response in this study. Poly(I:C) is a synthetic analog of dsRNA and mimics virus infection. Viruses have various strategies to antagonize the host defense system (*17*). However, poly(I:C) does not encode any virus-derived factors, therefore poly(I:C)-induced responses may mimic fundamental responses against virus infection. Here, we show that the regulatory networks of gene expression by TRBP through regulation of miRNA biogenesis during poly(I:C)-induced antiviral response.

## Results

### Poly(I:C) transfection induces antiviral response

During virus infection, virus-derived RNAs are sensed by RLRs and Toll-like receptor 3 (TLR3), resulting in production of type-I IFN. RLRs and TLR3 are different in subcellular localization: RLRs are localized in the cytoplasm, whereas TLR3 is localized on the cell membrane or in the endosome. We first confirmed which pathway is involved in IFN response induced by poly(I:C) transfection in human HeLa cells. IFN-β was significantly increased in mRNA level at 4 hours after poly(I:C) transfection (Fig. 1A), following the gradual increase in protein level (Fig. 1B). To determine IFN-β production pathway, we used specific siRNA (siIPS-1) to knock down IPS- 1, a key factor in the RLR pathway but not in the TLR3 pathway. The mRNA level of IFN-β was markedly decreased by siIPS-1 but not by negative control siRNA (siControl) during poly(I:C) transfection (Fig. 1C), suggesting that the transfected poly(I:C) was mainly sensed by RLRs in the cytoplasm and activated IFN response.

**Fig. 1:**
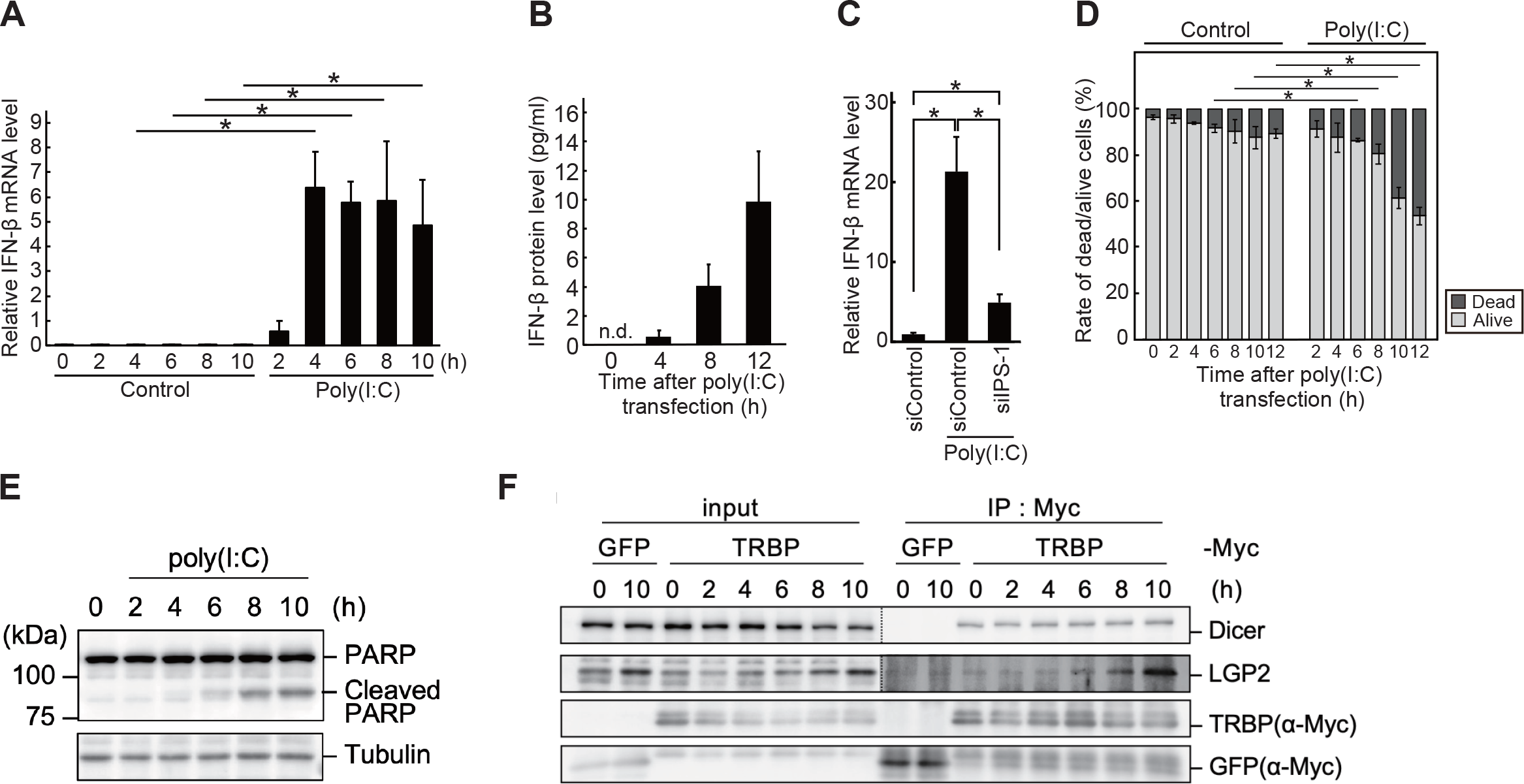
Poly(I:C) transfection induces antiviral responses. **(A)** Relative IFN-β mRNA level after poly(I:C) transfection was quantified by qRT-PCR, in the cells collected at 0, 2, 4, 6, 8, and 10 hours following mock treatment or poly(I:C) transfection. The experiments were performed in triplicate and P-values were determined by Student’ s t-test (*P < 0.05). **(B)** IFN-β protein level after poly(I:C) transfection was quantified by ELISA, using the culture medium of the cells collected at 0, 4, 8, and 12 hours following transfection. **(C)** Relative IFN-β mRNA level was quantified by qRT-PCR, using the cells transfected with or without siIPS-1 and collected at 8 hours following mock treatment or poly(I:C) transfection. The experiments were performed in triplicate and P-values were determined by Student’ s t-test (*P < 0.05). **(D)** Rate of dead and alive cells in WT cells after poly(I:C) transfection (n=4). The number of dead or alive cells was counted at 0, 2, 4, 6, 8, 10, and 12 hours following mock treatment or poly(I:C) transfection. P-values were determined by Student’ s t-test (*P < 0.05). **(E)** Western blot of endogenous PARP and tubulin in WT cells. The cells were collected at 0, 2, 4, 6, 8, and 10 hours following poly(I:C) transfection. **(F)** Western blot of immunoprecipitated LGP2 with TRBP. Plasmids encoding TRBP-Myc or GFP-Myc were transfected into HeLa cells, and the cells were collected at 0, 2, 4, 6, 8, and 10 hours following poly(I:C) transfection. Immunoprecipitation was performed with an anti-Myc antibody. Endogenous Dicer and LGP2 were detected by antibody against endogenous each protein.

Besides the IFN response, it is known that viral infection induces apoptotic cell death. We calculated the rate of dead cells every 2 hours until 10 hours after poly(I:C) transfection, and found that dead cells were significantly increased after 6 hours (Fig. 1D). Furthermore, in the apoptosis signaling pathway, the activated caspase-3 (CASP3) cleaves the 116 kDa form of poly (ADP-ribose) polymerase (PARP) to 85 and 31 kDa fragments (*18*). The results of western blotting showed that cleaved PARP was detected from 6 hours after poly(I:C) transfection (Fig. 1E), suggesting that transfected poly(I:C) induced apoptotic cell death.

Previously, we revealed that the upregulated LGP2 by SeV infection preferentially interacts with TRBP. Since LGP2 binds to TRBP through the dsRNA binding sites of pre-miRNA in TRBP, the amount of pre-miRNA binding to TRBP is competitively decreased (*10*, *14*). To examine whether LGP2 could also interact with TRBP in poly(I:C)-transfected cells, plasmids encoding C-terminal Myc-tagged TRBP (TRBP-Myc) and GFP (GFP-Myc) as a negative control were transfected into HeLa cells, respectively. Immunoprecipitation was performed using anti-Myc antibody at 0, 2, 4, 6, 8, and 10 hours after poly(I:C) transfection. TRBP and GFP were detected by anti-Myc antibody, and LGP2 was detected by antibody against endogenous LGP2 protein (Fig. 1F). LGP2 protein was obviously immunoprecipitated with TRBP at 6 hours after poly(I:C) transfection, suggesting that TRBP-LGP2 interaction is also induced by poly(I:C) transfection.

### Antiviral genes were induced by poly(I:C) transfection irrespective of TRBP

Next, we investigated the effect of TRBP on gene expression profiles in poly(I:C)-induced IFN response. Then, RNA-Sequencing (RNA-Seq) and small RNA-Seq were performed using wild-type (WT) and TRBP knockout (TRBP^-/-^) HeLa cells, generated in our previous reports (*14*), at 6, 8, and 10 hours after the poly(I:C) transfection (Fig. 2A). We identified upregulated genes (>2- fold) and downregulated genes (<0.5-fold) at 6 (Fig. S1A, B), 8 (Fig. S1C, D), 10 (Fig. 2B, C) hours after poly(I:C) transfection in WT and TRBP^-/-^ cells, respectively. IFN-β was induced at the highest levels at 6 hours after poly(I:C) transfection in WT and TRBP^-/-^ cells similarly determined by RNA-seq analyses (Fig. 2D). The amounts of upregulated genes were higher compared to those of downregulated genes in both WT and TRBP^-/-^ cells at any time points. Gene Ontology (GO) analyses of the upregulated genes revealed that the top four GO terms were completely the same in both cell types: transcription regulation, transcription, antiviral defense, and innate immunity (Fig. S1E-H, Fig. 2E, F). GO analyses of the downregulated genes showed that GO terms, lipid metabolism and transport, were identified in WT cells at any time points commonly, and GO terms, lipid metabolism, fatty acid metabolism, and electron transport, were identified in TRBP^-/-^ cells at any time points commonly (Fig. S1I-L, Fig. 2G, H). The genes included in antiviral defense or innate immunity in these GO terms were similar in WT and TRBP^-/-^ cells, whereas those in transcription regulation and transcription were not almost different. Thus, during poly(I:C) transfection, similar antiviral states were induced in WT and TRBP^-/-^ cells.

**Fig. 2:**
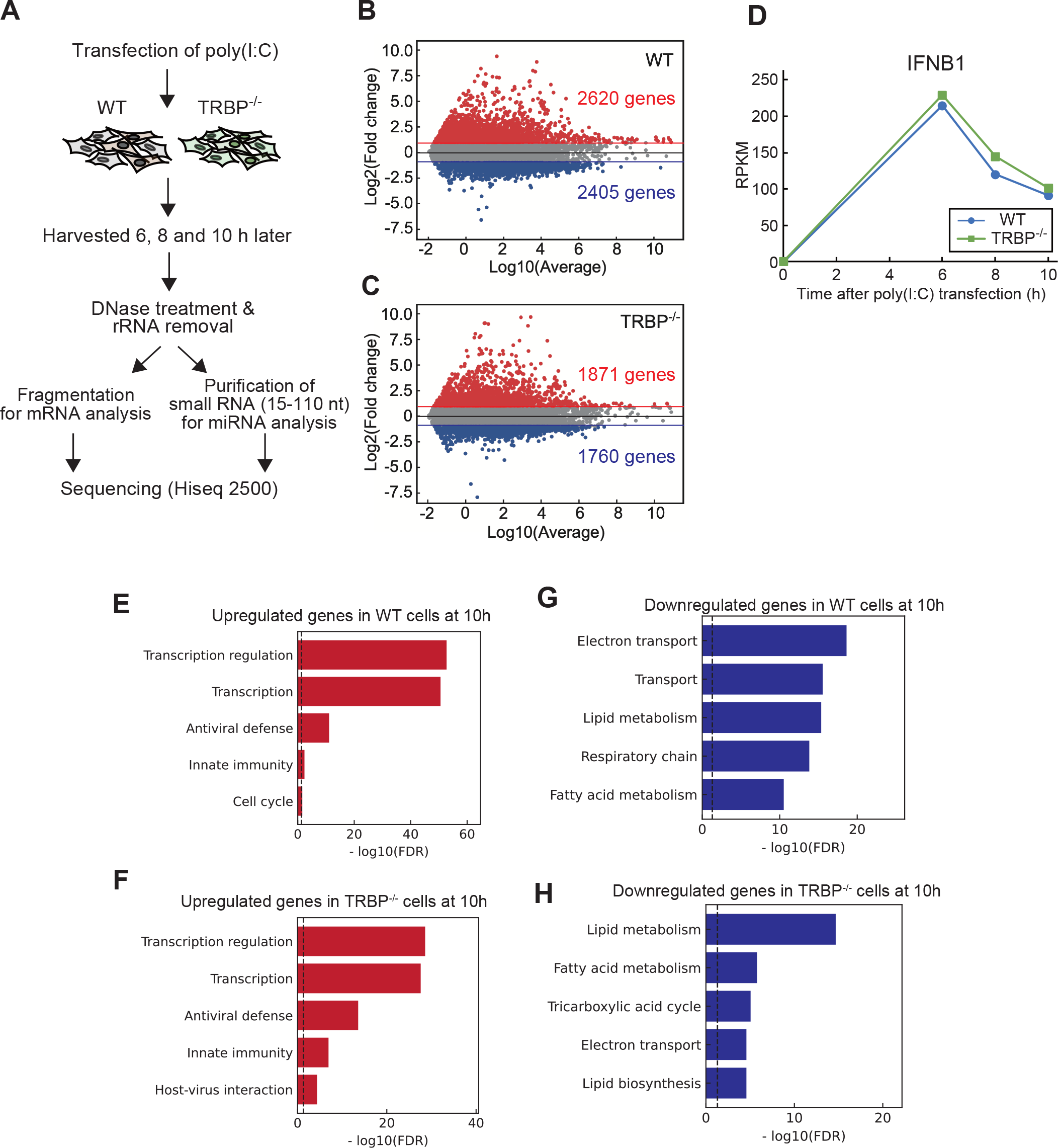
Changes in gene expression profiles by poly(I:C) transfection. (A) Schematic diagram of RNA-seq and small RNA-seq which were performed to analyze comprehensive gene expression profiles in WT or TRBP-/- cells after poly(I:C) transfection. **(B, C)** MA plot which shows changes of gene expression profile at 10 hours after poly(I:C) transfection in WT (B) or TRBP-/- (C) cells, respectively. A total of 15016 genes with RPKM > 0.1 were plotted. **(D)** Expression levels of IFN-β mRNA (IFNB1 gene) after poly(I:C) transfection were shown as RPKM calculated from RNA-seq results (n=2). **(E)** Top 5 GO terms enriched in upregulated genes at 10 hours after poly(I:C) transfection in WT cells. **(F)** Top 5 GO terms enriched in upregulated genes at 10 hours after poly (I:C) transfection in TRBP-/- cells. **(G)** Top 5 GO terms enriched in downregulated genes at 10 hours after poly(I:C) transfection in WT cells. **(H)** Top 5 GO terms enriched in downregulated genes at 10 hours after poly(I:C) transfection in TRBP-/- cells.

### Maturation of downregulated miRNAs by poly(I:C) transfection in WT cells may be regulated by TRBP

Previously, we discovered that IFN response suppressed the maturation of miRNAs by Dicer interacting with TRBP, since LGP2 upregulated by antiviral infection competitively interacts with TRBP. To confirm whether such suppression of miRNA maturation is observed in poly(I:C)- induced antiviral response, we performed RNA silencing activity assay as shown previously (*14*) (Fig. 3A). In this assay, we used miRNA mimic against firefly luciferase gene (miLuc), whose pre-miRNA (pre-miLuc) is bound to TRBP and cleaved into mature miLuc by Dicer, and downregulates the target luciferase activity. The poly(I:C) transfection significantly increased relative luciferase activity (Fig. 3B), meaning the repression of RNA silencing activity by the transfection of pre-miLuc, and the mature miLuc level was certainly downregulated by qRT-PCR (Fig. 3C). These results suggested that poly(I:C) transfection repressed maturation of TRBP- bound pre-miLuc by increasing TRBP-LGP2 interaction, and resulted in the decrease of RNA silencing activity by the decreased level of mature miLuc. Furthermore, to investigate the role of LGP2 in RNA silencing, LGP2 was knocked down by siRNA against LGP2 (siLGP2) and RNA silencing activity was measured (Fig. 3D). As a positive control, we investigated the effect of AGO2, which is an essential gene for RNA silencing. The AGO2 knockdown by siRNA against AGO2 (siAGO2) increased luciferase activity as expected, but the LGP2 knockdown significantly decreased luciferase activity. Thus, our results indicated that LGP2 suppressed RNA silencing activity in poly(I:C)-transfected cells.

**Fig. 3:**
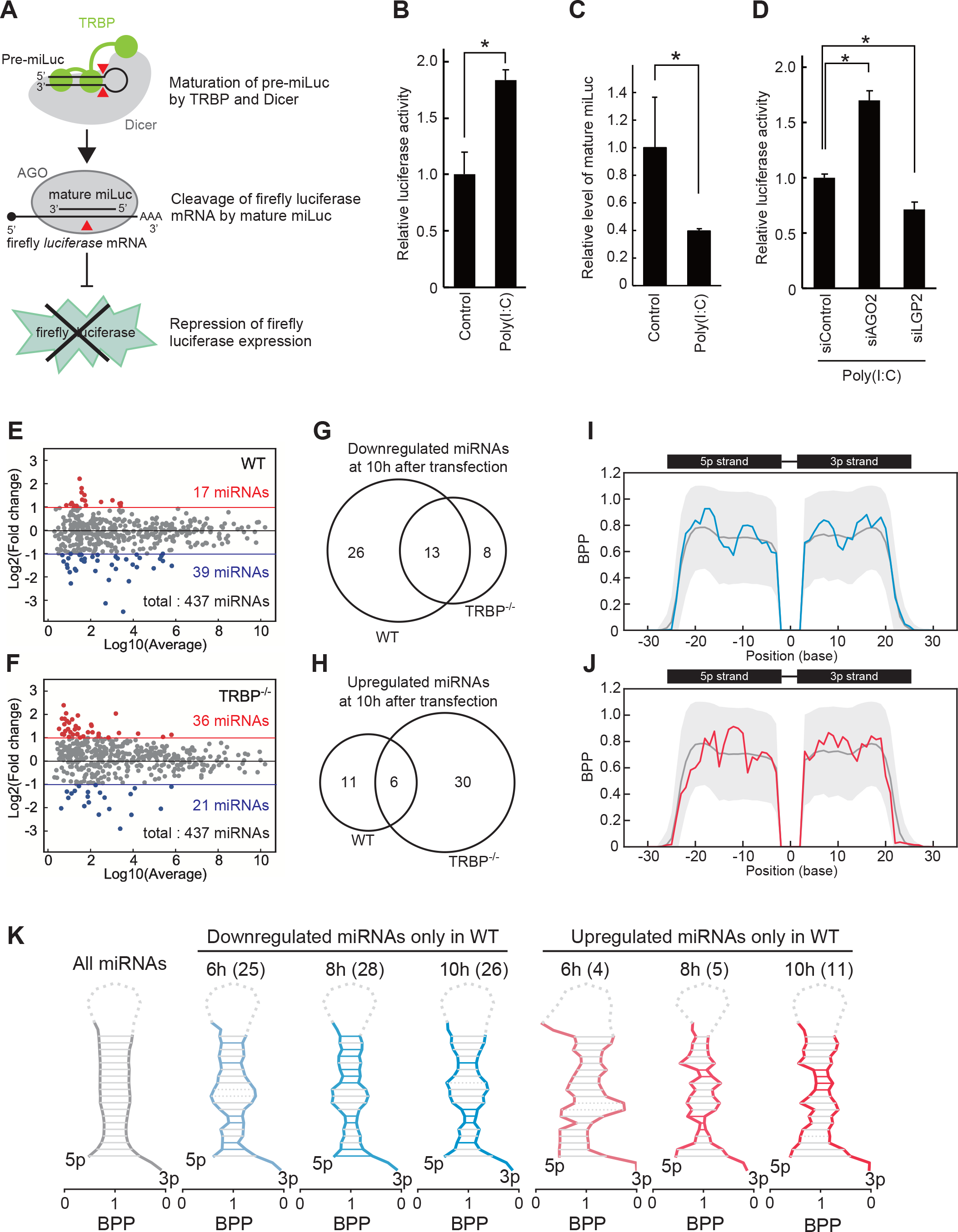
Changes in expression profiles of mature miRNAs by poly(I:C) transfection. (A) Schematic diagram of reporter assays. RNA silencing activities were measured by the luminescence of firefly luciferase which is the target of mature miLuc. **(B)** Luciferase reporter assay of the cells transfected with pre-miLuc during poly(I:C) transfection. After transfection of pre-miRNA, the cells were collected at 24 hours after mock treatment or poly(I:C) transfection. The experiments were performed in triplicate and P-values were determined by Student’ s t-test (*P < 0.05). **(C)** Relative level of mature miLuc at 24 hours after poly(I:C) transfection was quantified by qRT-PCR. The experiments were performed in triplicate and P-values were determined by Student’ s t-test (*P < 0.05). **(D)** Luciferase reporter assay of the cells transfected with siControl, siAGO2 and siLGP2, and collected at 24 hours after poly(I:C) transfection. The experiments were performed in triplicate and P-values were determined by Student’ s t-test (*P < 0.05). **(E, F)** MA plots which indicate changes of mature miRNA expression levels at 10 hours after poly(I:C) transfection in WT (E) and TRBP-/- (F) cells, respectively. **(G, H)** Venn’ s diagrams which show overlaps between downregulated (G) and upregulated (H) miRNAs in WT and TRBP-/- cells at 10 hours after poly(I:C) transfection. **(I, J)** BPP values in the registered pre-miRNA structures of downregulated (I) or upregulated (J) mature miRNAs were calculated using the max value of base-pairing probabilities calculated by CentroidFold. The gray line indicates the mean BPP at each position of all pre-miRNAs registered in miRBase, and the gray area indicates the standard deviation. The average BPP of miRNAs which was downregulated only in WT cells was shown in blue (I) and those upregulated only in WT cells were shown in red (J) at 10 hours after poly(I:C) transfection. **(K)** Predicted secondary structure of all pre-miRNAs (gray), downregulated miRNAs (blue) and upregulated miRNAs (red) only in WT cells reconstructed using BPP values. The stem-loop structure of pre-miRNA was created from a 5p strand and a 3p strand sequences with two nucleotide overhangs. Solid lines of blue or red show the positions with significantly higher BPPs determined by Student’ s t-test (*P < 0.05).

Then, we examined comprehensive effects of poly(I:C) transfection on endogenous miRNAs. The changes of the expression profiles of mature miRNAs by poly(I:C) transfection were examined by small RNA-Seq analysis. To determine the miRNA expression level, reads per million mapped reads (RPM) were calculated for each mature miRNA registered in miRbase (*19–21*). We identified 6, 9, and 17 upregulated mature miRNAs (>2-fold), and 39, 45, and 39 downregulated mature miRNAs (<0.5-fold) at 6 (Fig. S2A), 8 (Fig. S2E), and 10 (Fig. 3E) hours after poly(I:C) transfection in WT cells, respectively. On the other hand, 44, 43, and 36 upregulated miRNAs and 18, 28, and 21 downregulated miRNAs were detected at 6 (Fig. S2B), 8 (Fig. S2F), and 10 (Fig. 3F) hours in TRBP^-/-^ cells, respectively. To clarify the essential impact of TRBP on miRNA expression profiles, we focused on the upregulated or downregulated miRNAs only in WT cells but not in TRBP^-/-^ cells. There were 25 (Fig. S2C), 28 (Fig. S2G), 26 (Fig. 3G) downregulated miRNAs and 4 (Fig. S2D), 5 (Fig. S2H), 11 (Fig. 3H) upregulated miRNAs only in WT cells at 6, 8, and 10 hours, respectively. Thus, the expression of many types of mature miRNAs was considered to be downregulated by poly(I:C) transfection by means of TRBP. To compare the features of these miRNAs, the secondary structures of pre-miRNAs of these mature miRNAs were analyzed by calculating base-pairing probabilities (BPPs), since our previous report suggested that TRBP-bound miRNAs have characteristic structural features (*14*). The structures of miRNAs downregulated only in WT cells exhibited specific features with high BPPs in the stem regions except for a central region of pre-miRNAs at any time point (Fig. 3I, K, S3A, C). However, such distinctive structures were not observed in miRNAs upregulated only in WT cells (Fig. 3J, K, S3B, D) or those downregulated and upregulated miRNAs in TRBP^-/-^ cells (Fig. S3E). These results strongly suggested that the maturation of TRBP-bound miRNAs was suppressed in WT cells by poly(I:C) transfection, probably because miRNA maturation by TRBP is suppressed by LGP2 induction by poly(I:C).

As a cause of the differential levels of mature miRNAs after poly(I:C) transfection, there is a possibility that the expression levels of pri-miRNAs were changed and the maturation steps of pri- or pre-miRNAs were not regulated. Then, we focused on the miRNAs transcribed from the host genes, since the expression levels of such miRNAs are correlated with the expression levels of host genes (*22*, *23*). At first, we investigated the genomic locations of all of 2573 miRNAs registered in miRbase and identified their host genes from the miRAID database (*24*). The results indicated that 65.2% of miRNAs were located in the intragenic region, and 34.8% of them were in the intergenic region (Fig. S4A). The genomic locations of 446 miRNAs expressed in HeLa cells also showed similar results (63.8% were located in the intragenic region and 36.2% were in the intergenic region) (Fig. S4A). The log2 fold changes (FCs) of reads of these intragenic miRNAs in WT cells during poly(I:C) response might be slightly decreased compared to those in TRBP^-/-^ (Fig. S4B). However, log2 FCs of the reads of their host genes were almost similar in both WT and TRBP^-/-^ cells (Fig. S4C). The relationship between the FCs of intragenic miRNAs and their host genes were analyzed by calculating correlation coeffieints (Fig. S4D-I). As a result, there were no clear correlations between the FCs of intragenic miRNAs and their host genes, suggesting that the expression levels of host genes did not affect the mature miRNA levels during poly(I:C) response. Thus, we suggest that in the step of pre-miRNA processing by TRBP is a major reason in a decrease in the mature miRNA levels.

### Predicted regulatory network of gene expression by downregulated mature miRNAs in WT cells during poly(I:C) transfection

To uncover the overall picture of gene expression network regulated by the TRBP-bound miRNAs whose maturation is downregulated by poly(I:C) transfection, we predicted the target genes of the downregulated miRNAs using TargetScan (*25*) (Fig. 4A), and the functions of predicted target genes were determined by GO analysis using DAVID web server (*26*, *27*). To refine the functional meaning of TRBP-regulated miRNAs which were considered to be downregulated only in WT cells at 10 hours after poly(I:C) transfection (Fig. 4B), the target genes of downregulated miRNAs only in WT cells and others (miRNAs downregulated in TRBP^-/-^ cells) were predicted. From such genes, the genes actually upregulated by poly(I:C) transfection were examined by GO analysis. Interestingly, GO analysis clearly revealed that transcription-related genes categorized into GO terms including transcription regulation, and transcription were highly enriched as the predicted targets of the downregulated miRNAs in either WT or TRBP^-/-^ cells (Fig. 4C-E). Similar results were also observed for miRNAs downregulated at 6 (Fig. S5A-D) or 8 (Fig. S5I-K) hours after poly(I:C) transfection. However, the genes in these transcription-related genes predicted as target genes of the downregulated miRNAs were different in WT or TRBP^-/-^ cells (4B, S5A, H). Then, we next examined the downstream target genes of these transcription-related genes in either WT or TRBP^-/-^ cells or both. The predicted target genes of downregulated miRNAs only in WT cells, actually upregulated (>2-fold) transcription factors were listed up at 6, 8, or 10 hours after poly(I:C) transfection, and their target genes were predicted using TFlink database (*28*). The GO analyses against these target genes revealed that the genes involved in the GO terms, cell cycle and apoptosis, were most abundantly detected at any time points after poly(I:C) transfection (Fig. 4F, S5E, L). Such genes were not predicted to be the targets of transcription-regulated genes upregulated in TRBP^-/-^ cells or both WT and TRBP^-/-^ cells (Fig. 4G, H, S5F, G, M, N). Our previous report also revealed that apoptosis-related genes are regulated by TRBP-LGP2 interaction. Therefore, it was strongly suggested that the expression of apoptosis-related genes is regulated directly by downregulated miRNAs and indirectly through transcription-related genes in WT cells by poly(I:C). Thus, such gene expression network is considered to be regulated by TRBP. The detailed gene regulatory network, orchestrated by miRNAs and transcription factors, was created, especially extracting the regulatory networks starting from the downregulated miRNAs in WT cells by poly(I:C) and ending in apoptosis- or cell cycle-related genes (Fig. 4I, Table2, 3).

**Fig. 4:**
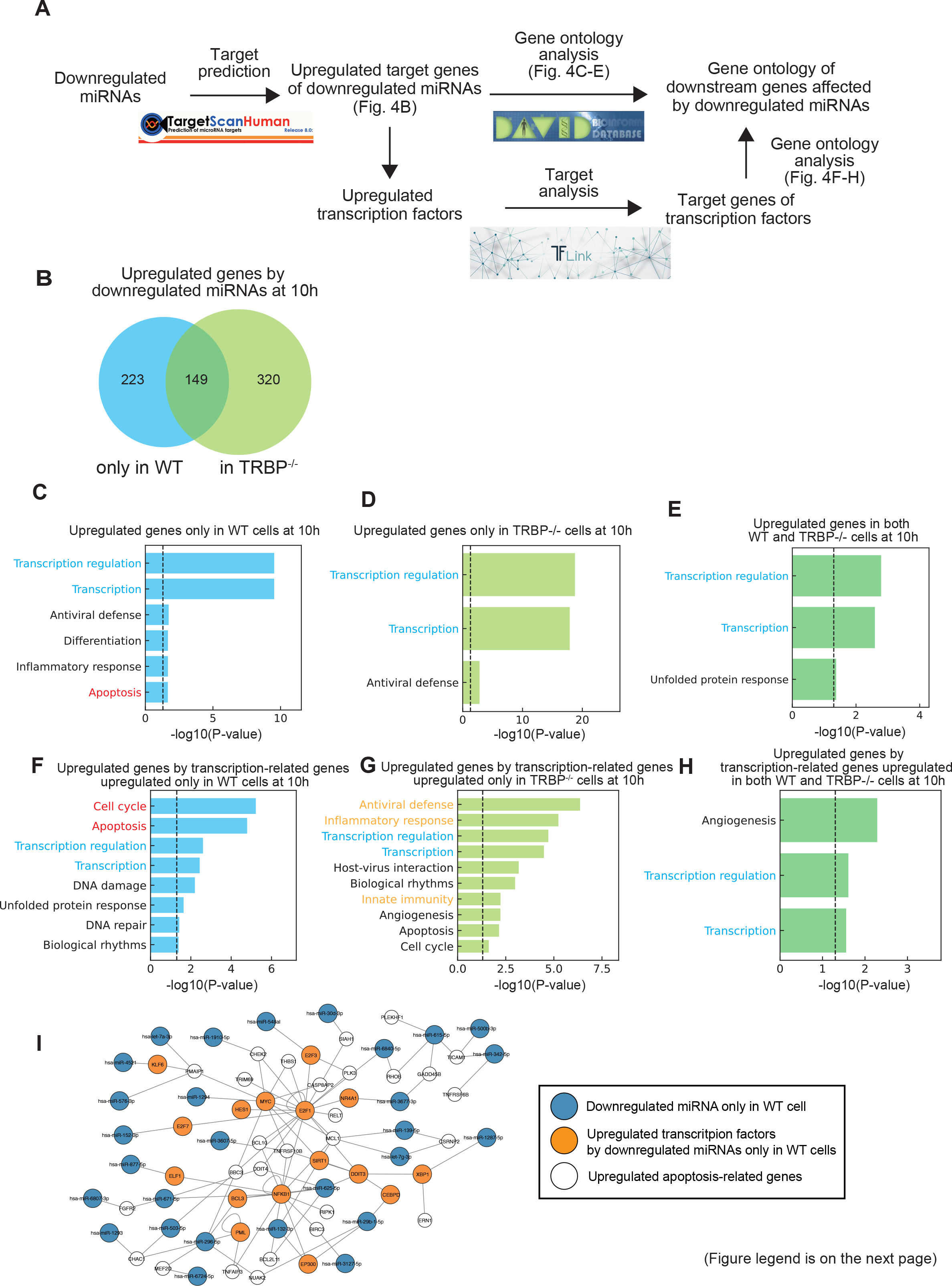
Predicted regulatory network of gene expression by downregulated mature miRNAs in WT cells during poly(I:C) transfection. (A) Schematic diagram of target prediction analyses using TargetScan, TFlink databases and DAVID web server. **(B)** Venn’ s diagram indicating the overlap between upregulated genes by downregulated miRNAs in only WT cells and in TRBP-/- cells at 10 hours after poly(I:C) transfection. **(C)** Enriched GO terms in upregulated genes only in WT cells at 10 hours after poly(I:C) transfection (sky-blue region in Fig. 4B). **(D)** Enriched GO terms in upregulated genes only in TRBP-/- cells at 10 hours after poly(I:C) transfection (light-green region in Fig. 4B). **(E)** Enriched GO terms in upregulated genes both in WT and TRBP-/- cells at 10 hours after poly(I:C) transfection (green region in Fig. 4B). **(F)** Enriched GO terms in upregulated target genes by transcription-related genes, specifically upregulated only in WT cells at 10 hours (Term “Transcription” in Fig. 4C). **(G)** Enriched GO terms in upregulated target genes by transcription factors, specifically upregulated only in TRBP-/- cells at 10 hours (Term “Transcription” in Fig. 4D). **(H)** Enriched GO terms in upregulated target genes by transcription factors, commonly upregulated both in WT and TRBP-/- cells at 10 hours (Term “Transcription” in Fig. 4E). **(I)** The regulatory network of upregulated apoptosis-related genes by downregulated miRNAs only in WT cells. In the result of GO analysis, genes which were categorized in Term “Apoptosis” were extracted as apoptosis-related genes and used for this network.

### Validation of miRNA-mRNA network of apoptosis induced by poly(I:C)

The predicted miRNA-mRNA network using target prediction databases was constructed in Fig. 4I. To investigate whether such a network is accurate or not, we experimentally examined the upstream and downstream gene regulatory relationships. Among the apoptosis-related genes which were predicted as downregulated miRNA targets in WT cells but not in TRBP^-/-^ cells, there were well-known pro-apoptotic genes such as Phorbol-12-Myristate-13-Acetate-Induced Protein 1 (PMAIP1) (also known as Noxa) and BCL2 like 11 (BCL2L11) (also known as Bim) (Fig. 4I, Table 2). PMAIP1 and BCL2L11 are members of BH3-only BCL-2 family proteins, and known as pro-apoptotic genes which are essential initiators of apoptosis. During poly(I:C) transfection, the mRNA levels of these pro-apoptotic genes were revealed to be highly upregulated in WT cells compared to TRBP^-/-^ and LGP2^-/-^ cells by determined by qRT-PCR (Fig. 5A, B). Their expression levels were only slightly upregulated in TRBP^-/-^ and LGP2^-/-^ cells (Fig. 5A, B), suggesting that TRBP and LGP2 are necessary to upregulate the expression of these genes. Thus, these results strongly suggest that the upregulated LGP2 by poly(I:C) transfection interacted with TRBP, resulting in the suppression of miRNA maturation by Dicer.

**Fig. 5:**
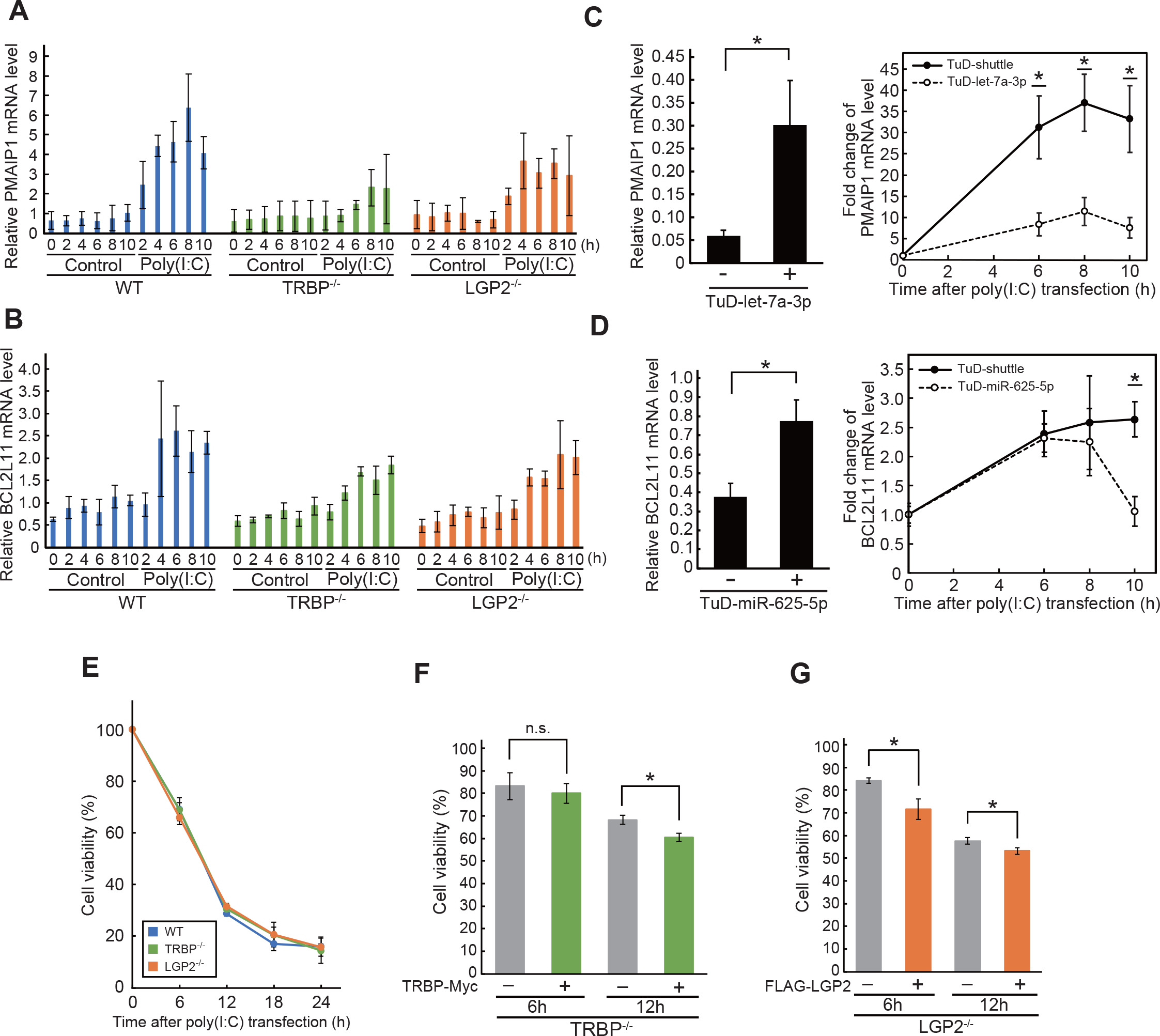
Validation of miRNA-mRNA network of apoptosis induced by poly(I:C). (A, B) Relative PMAIP1 (A) or BCL2L11 (B) mRNA level after poly(I:C) transfection was quantified by qRT-PCR. WT, TRBP-/- and LGP2-/- HeLa cells were collected at 0, 2, 4, 6, 8, and 10 hours following mock treatment or poly(I:C) transfection. The experiments were performed in triplicate and P-values were determined by Student’ s t-test (*P < 0.05). **(C, D)** Relative PMAIP1 (C) or BCL2L11 (D) mRNA level with TuD miRNA inhibitor was quantified by qRT-PCR. WT HeLa cells were transfected TuD-shuttle control vector or TuD-let-7a-3p or TuD-miR-625-5p one day before poly(I:C) transfection. Cells were collected at 0, 6, 8, and 10 hours following poly(I:C). PMAIP1 mRNA levels at 0 hours were shown in the left panel, and time-dependent changes in expression were shown in the right panel. The experiments were performed in triplicate and P-values were determined by Student’ s t-test (*P < 0.05). **(E)** Cell viability of WT, TRBP-/- and LGP2-/- HeLa cells after poly(I:C) transfection. The number of alive cells was counted at 6, 12, 18, and 24 hours following mock treatment or poly(I:C) transfection. Cell viability was calculated by this formula: (The number of alive cells in poly(I:C)-transfected sample) / (The number of alive cells in mock-treated sample). **(F, G)** Cell viability of TRBP-/- (F) or LGP2-/- (G) HeLa cells after poly(I:C) transfection with or without overexpression of TRBP-Myc. The number of alive cells was counted at 6 and 12 hours following mock treatment or poly(I:C) transfection. Cell viability was calculated by following formula: (The number of alive cells in poly(I:C)-transfected sample) / (The number of alive cells in mock-treated sample).

**Table 1.**
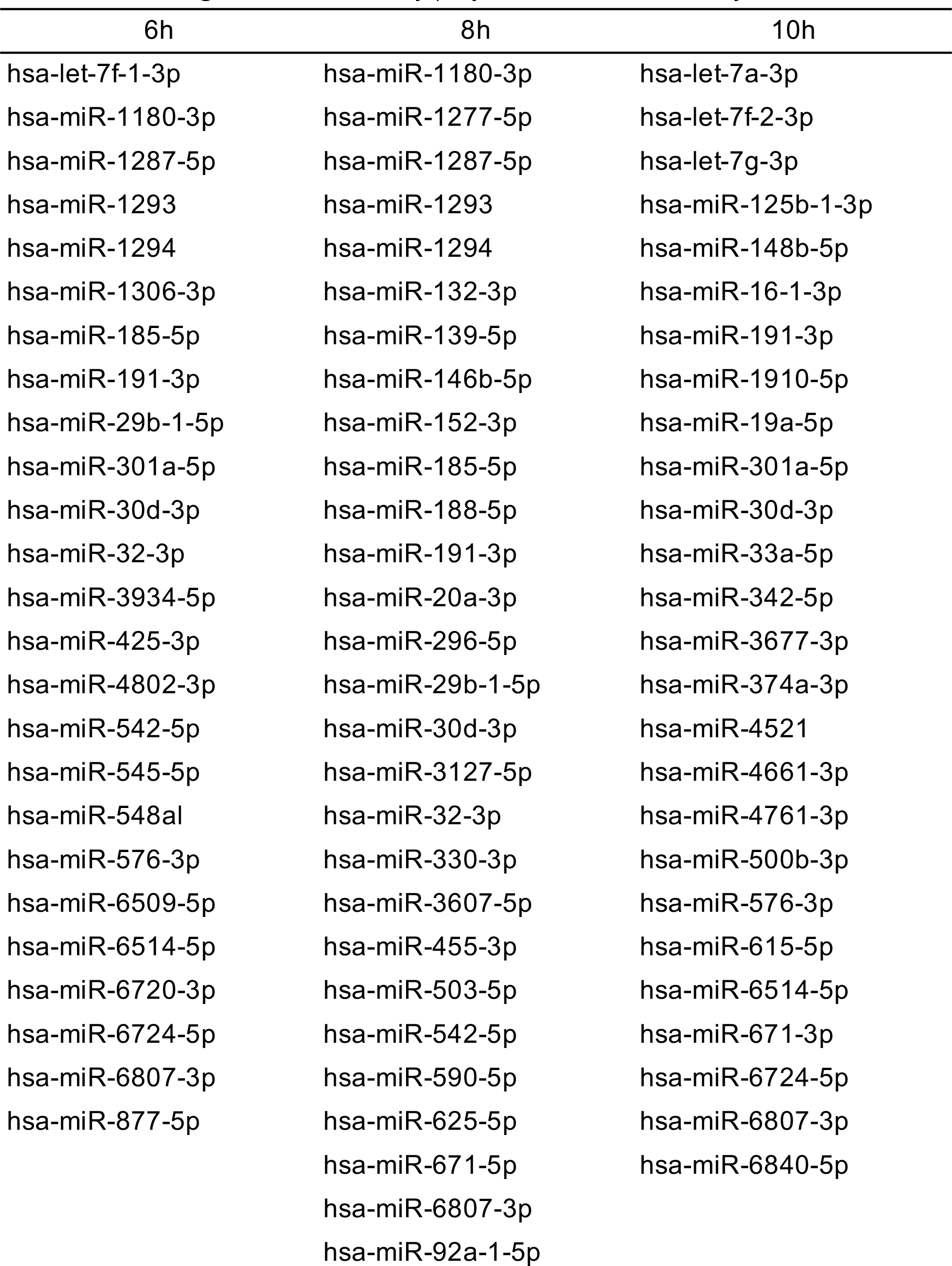
Downregulated miRNAs by poly(I:C) transfection only in WT cell.

**Table 2.**
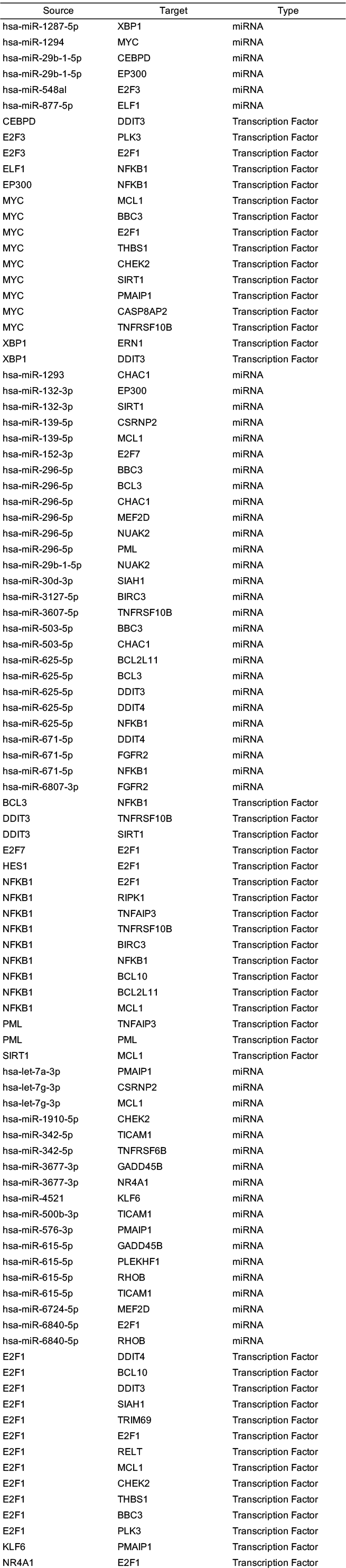
Predicted regulatory network of upregulated apoptosis-related genes from miRNAs and transcription factors.

**Table 3.**
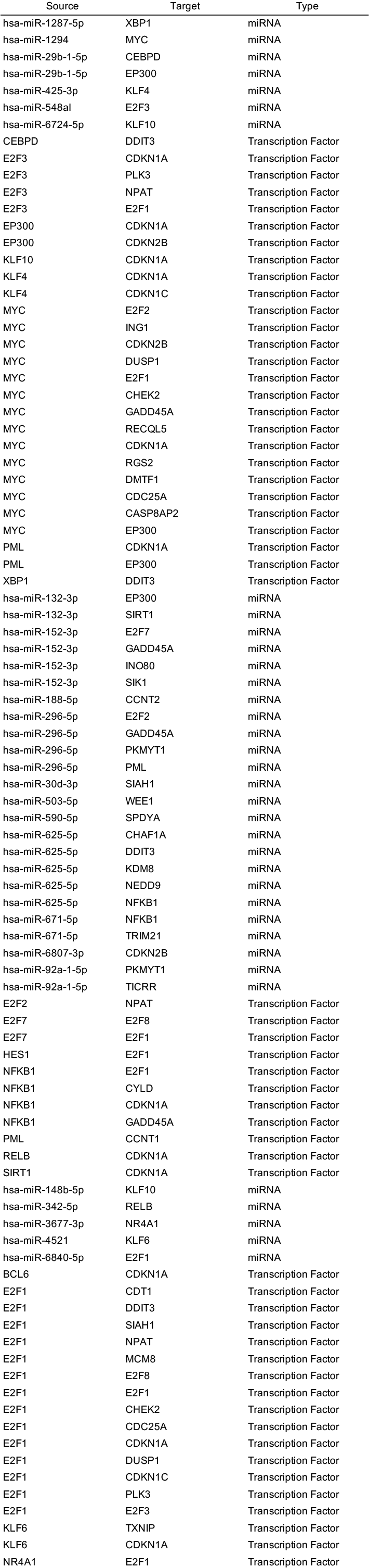
Predicted regulatory network of upregulated cell cycle-related genes from miRNAs and transcription factors.

In this study, PMAIP1 was predicted to be a target gene of let-7a-3p, and BCL2L11 is a target of miR-625-5p, and both miRNAs were downregulated only in WT cells but not in TRBP^-/-^ cells (Fig. S6A, B).

To validate the effects of these miRNAs on the expression of target genes, we inhibited the functions of miRNAs by Tough Decoy miRNA inhibitor (TuD), which has miRNA-binding sites with complementarity sequence specific to each miRNA (*29*). The plasmid encoding a specific TuD inhibitor for let-7a-3p (TuD-let-7a-3p) was transfected into WT HeLa cells and PMAIP1 mRNA expression was detected by qRT-PCR (Fig. 5C). The introduction of TuD-let-7a-3p significantly increased PMAIP1 mRNA levels, suggesting that TuD-let-7a-3p inhibited the function of let-7a-3p which suppressed PMAIP1 expression in WT HeLa cells. Furthermore, the upregulation of PMAIP1 mRNA level during poly(I:C) treatment was apparently repressed in the presence of TuD-let-7a-3p, compared to the cells treated with TuD-shuttle control vector. From these results, it was indicated that PMAIP1 is certainly regulated by let-7a-3p as a target gene, and the inhibition of let-7a-3p maturation during poly(I:C) transfection should be essential in the induction of PAMIP1 mRNA. We also examined the effect of miR-625-5p on BCL2L11 expression, using TuD-miR-625-5p. BCL2L11 was significantly upregulated after transfection of TuD-miR-625-5p (Fig. 5D), indicating that BCL2L11 expression is suppressed by miR-625-5p in the normal condition. As for the changes in the expression levels of BCL2L11 during poly(I:C) treatment, the degree of induction was not affected by transfection of TuD-miR-625-5p. However, miRNA levels of BCL2L11 peaked out rapidly after 8h from poly(I:C) transfection in the presence of TuD-miR-625-5p, suggesting that the regulation of BCL2L11 expression by miR-625-5p is essential for prolonged upregulation of BCL2L11 during the poly(I:C) transfection, or such miR- 625-5p-BCL2L11 regulation may necessary after 8 hours.

Next, we investigated whether the expression of pro-apoptotic genes induces cell death which is also induced by poly(I:C) transfection. Poly(I:C) were transfected into WT, TRBP^-/-^, and LGP2^-/-^ HeLa cells and surviving cells were counted after 6, 12, 18 and 24 hours (Fig. 5E). The cell viability of WT cells did not show any significant differences with those of TRBP^-/-^ or LGP2^-/-^ cells, but was slightly lower at 12 and 18 hours. When TRBP-Myc construct was transfected into TRBP^-/-^ cells or Flag-LGP2 construct transfected into LGP2^-/-^ cells, cell viabilities were significantly decreased after 12 hours of poly(I:C) transfection (Fig. 5F, G). Therefore, it was confirmed that either TRBP or LGP2 related to the induction of cell death during poly(I:C) transfection, probably through upregulation of pro-apoptotic genes by the inhibition of miRNA maturation.

## Discussion

In this report, we confirmed that poly(I:C) transfection induces a similar condition of IFN response by viral infection. The mRNA of IFN was induced at first (Fig. 1A), and followed by the increase of its protein level (Fig. 1B). Furthermore, apoptosis verified by cell death induction and PARP cleavage was also induced (Fig. 1D, E). Furthermore, LGP2, an ISG protein, was also induced (Fig. 1F). Thus, poly(I:C) transfection is considered to induce the responses which mimic fundamental responses against virus infection without encoding any virus-derived factors. The effect of poly(I:C) transfection on RNAi activity was determined by reporter assay using miLuc (Fig. 3A). RNAi activity was repressed by poly(I:C) transfection (Fig. 3B), and it was recovered by knockdown of LGP2 (Fig. 3D). LGP2 is a TRBP-interacting protein, and one of ISGs which are upregulated during IFN response. Therefore, it was suggested that pre-miLuc maturation was inhibited by upregulation of LGP2 expression upon poly(I:C) transfection via TRBP-LGP2 interaction, since TRBP-LGP2 interaction competitively inhibits the TRBP-Dicer interaction which enhances pre-miLuc maturation.

We previously discovered that TRBP-bound miRNAs regulate apoptosis-related genes during SeV infection, but a comprehensive analysis of miRNA expression and their regulation of downstream mRNA expression was not conducted (*10*). In this report, we revealed that TRBP- bound miRNAs play a role in inducing apoptosis in antiviral response. We performed RNA-Seq and small RNA-Seq to analyze the differentially expressed endogenous miRNAs as well as mRNAs. Analysis of miRNAs revealed that the number of decreased miRNAs in WT cells alone was apparently larger compared to those in TRBP^-/-^ cells at any time points after poly(I:C) transfection (Fig. 3G, S2C, G), suggesting that these miRNAs are considered to be downregulated by the decreased interaction of TRBP-Dicer by the upregulation of LGP2 expression. Furthermore, these WT-specifically downregulated miRNAs showed characteristic secondary structures for pre-miRNAs for TRBP binding (Fig. 3I, K).

These pre-miRNAs showed weak BPPs in the center of the miRNA duplexes and high BPPs in both sides of the center (Fig. 3K). In contrast, such clear characteristics were not observed in the miRNA duplexes upregulated only in WT cells (Fig. 3K). TRBP has three dsRBDs, of which dsRBD1 and dsRBD2 bind to miRNA duplexes, but dsRBD3 binds to Dicer. In the processing step by the TRBP-Dicer complex, processing efficiency is higher when there are fewer mismatches near the cleavage point (*30*). Thus two dsRBDs are thought to bind to regions of strong base pairing. Therefore, two regions with high BPPs in miRNA duplexes may be necessary for TRBP binding, and 4 RNA base-pairs were shown to be necessary for TRBP binding (*31*). Thus two TRBP binding positions should be situated at both sides of the central regions of miRNA duplexes.

In contrast to the downregulated mature miRNAs in WT cells alone, there are also downregulated miRNAs in TRBP^-/-^ cells or both WT and TRBP^-/-^ cells, and the upregulated miRNAs in either or both WT and TRBP^-/-^ cells. The maturation of these miRNAs is unlikely regulated by TRBP. Therefore, the expression levels of their host genes were examined (Fig. S4). As a result, the expression levels of host genes showed no effects on the expression levels of mature miRNAs. Then, there are possibilities that their maturation might be caused by some other dsRBPs, such as protein activator of interferon-induced protein kinase EIF2AK2 (PRKRA)(also known as PACT) and adenosine deaminase RNA specific (ADAR). Especially, ADAR1 gene has an IFN-induced isoform called ADAR1 p150, therefore ADAR1 function on miRNA biogenesis which is usually covered by TRBP function might have contributed to the upregulation of mature miRNAs in TRBP^-/-^ cells. The upregulated genes predicted to be the targets of downregulated miRNAs in WT or TRBP^-/-^ cells after poly(I:C) transfection were categorized into similar GO terms of transcription regulation, and transcription (Fig. 4C, D, E, S5B, C, D, I, J, K). Their predicted target genes were apparently different: the predicted targets of transcription-related genes of WT cells were cell cycle and apoptosis (Fig. 4F, S5E, L), those of TRBP^-/-^ cells were antivirus defense or inflammatory response (Fig. 4G, S5F, M). The results suggested that the upregulated transcription-related factors were differentially regulated by types of miRNAs, and such types of miRNAs are considered to be certainly regulated by the processing steps regulated by TRBP. Thus, TRBP may function as the regulator of the gene expression network during IFN response.

When poly(I:C) induces apoptosis, cell cycle-related genes were also upregulated. Kim et al. revealed that phosphorylation of TRBP is closely related to cell cycle regulation through TRBP- PKR interaction (*32*). Analysis of this report suggests that not only phosphorylation and protein-protein interactions but also miRNA-mediated gene regulation might affect cell cycles. At the same time, DNA fragmentation which occurs during apoptosis is known to induce cell cycle arrest via activation of p53 (*33*). Therefore, it might be possible that genes involved in the cell cycle were affected through DNA damage in the apoptotic processes.

## Materials and methods

### Cell culture

Human WT, TRBP^-/-^, and LGP2^-/-^ HeLa cells were cultured in Dulbecco’s Modified Eagle’s Medium (Wako) with 10% Fetal Bovine Serum (Gibco) at 37°C with 5% CO2. TRBP^-/-^ and LGP2^-/-^ HeLa cells were generated using CRISPR/Cas9 system in our previous report (*14*).

### siRNA

The siRNAs used for gene knockdown experiments were chemically synthesized (Sigma-Aldrich, Shanghai GenePharma). The sequences of siRNAs are shown in Table S1.

### Plasmid construction

To construct the expression plasmid of C-terminal Myc-tagged TRBP (TRBP-Myc), a full-length human TRBP was amplified by PCR using cDNA synthesized from total RNA of HeLa cells, and cloned into pcDNA3.1-myc/His A vector (Invitrogen) with restriction enzymes, HindIII and NotI, as described previously (*34*). For the expression plasmid of N-terminal Flag-tagged LGP2 (Flag- LGP2), a full-length human LGP2 was amplified by PCR using cDNA synthesized from total RNA of HeLa cells, and cloned into the pcDNA3.1 vector. A sequence encoding the Flag tag was inserted at the N-terminal side of LGP2.

For the construction of a plasmid expressing inhibitor of miRNAs, named Tough Decoy (TuD), oligonucleotide pairs were annealed and cloned into the pmU6-TuD-shuttle digested with BsmBI. The sequences of oligonucleotides were shown in Table S2.

### Dual-luciferase reporter assay

RNA silencing activity was measured by luciferase reporter assay. One day before transfection, WT HeLa cells or those transfected with each siRNA were plated at 1 x 10^5^ cells / well in 24-well plate. Cells were transfected with 0.5 μg of pGL3-Control vector (Promega) encoding firefly luciferase, 0.5 μg of pSilencer-3.1-H1-puro vector encoding stem-loop-structured RNA resembling pre-miRNA targeting firefly luciferase (pre-miLuc), 0.1μg of pRL-SV40 vector (Promega) encoding *Renilla* luciferase using Lipofectamine 2000 (Invitrogen). A day after transfection, cells were transfected with poly(I:C) at 0.5 µg / well using Lipofectamine 2000. After 24 hours, cells were lysed with 100 μL of 1 x PLB (Promega). Luciferase activities were measured by Dual-Luciferase Reporter Assay System (Promega), and relative luciferase activity was calculated by the following formula; (firefly luciferase activity) / (Renilla luciferase activity).

### qRT-PCR

One day before transfection, WT HeLa cells were plated at 1 x 10^5^ cells / well in a 24-well plate. The cells were transfected with poly(I:C) (*12*) at 1 µg/well using Lipofectamine 2000. For inhibiting specific miRNA, a plasmid encoding TuD miRNA inhibitor was transfected into the cells a day before the transfection of poly(I:C). For RT-PCR, the cells treated with poly(I:C) were harvested at every 2 h until 10 h after the transfection. In the case of poly(I:C) treatment without transfection, poly(I:C) was added to the culture medium at 1 µg/well. Total RNAs were extracted from the cells using ISOGEN (NIPPON GENE). The cDNA was synthesized using High Capacity cDNA Reverse Transcription Kits (Applied Biosystems). For reverse transcription, random primers supplied with the kit were used for mRNA, and stem-loop primers were used for reverse transcription of mature miLuc. mRNA level of each gene was measured by quantitative RT-PCR (qRT-PCR) using the cDNAs as templates. As a control, mRNA level of tubulin was measured. The primers used for RT-PCR are shown in Table S3.

### ELISA assay

WT HeLa cells were plated at 1 x 10^5^ cells / well in a 24-well plate before 24 hours of poly(I:C) transfection. After poly(I:C) transfection, cell growth medium was collected every 4 h until 12 h after poly(I:C) transfection. Using the 50 µL of collected medium, the concentration of IFN-β protein in the medium was quantified by ELISA kit (VeriKineTM - HS Human IFN-β TCM ELISA Kit, PBL Assay Science).

### Cell viability assay

WT HeLa cells were plated at 1 x 10^5^ cells / well in a 24-well plate before 24 hours of poly(I:C) transfection. After poly(I:C) transfection, cells were collected in Trypsin-EDTA (Wako) and stained with 0.4% (w/v) Trypan Blue Solution (Wako) at each 6 hours. Unstained cells were counted and cell viabilities of poly(I:C)-transfected cells normalized to those of mock-treated cells were calculated.

### Immunoprecipitation assay

One day before transfection, WT HeLa cells were plated at 1.2 x 10^6^ cells / dish in a 6 cm dish. The cells were transfected with a plasmid encoding TRBP-Myc or GFP-Myc as a negative control using polyethylenimine (PEI). One day later, the cells were transfected with poly(I:C) with Lipofectamine 2000 and harvested at the indicated time. The cells were washed with 1 x PBS and lysed in lysis buffer (10 mM Hepes-NaOH (pH 7.9), 1.5 mM MgCl2, 10 mM KCl, 0.5 mM DTT, 140 mM NaCl, 1 mM EDTA, 1 mM Na3VO4, 10 mM NaF, 0.5% NP-40 and complete protease inhibitor). The cell lysates were mixed with 30 µL of Protein G Sepharose 4B (Sigma-Aldrich) and rotated at 4°C for 2 h with 2.5 µg of mouse anti-Myc antibody (Calbiochem). The beads were washed twice with wash buffer containing 300 mM NaCl and once with lysis buffer. To elute the bound proteins, 2 x SDS-PAGE sample buffer (30 µL) was added and the beads were boiled for 5 min.

### Western blotting

The eluted proteins from beads were separated by 10% (for detection of RLRs and Dicer) or 12.5 % (for detection of TRBP-Myc and GFP-Myc) polyacrylamide gel electrophoresis and western blotting, and transferred on polyvinylidene fluoride membranes by the Trans-Blot Turbo Transfer System (Bio-Rad). The membranes were treated in the blocking buffer, TBS-Tween (TBS-T; 20 mM Tris-HCl (pH 7.5), 150 mM NaCl, 0.1% Tween 20) with 5% Difco skim milk (Becton, Dickinson and Company), for 1 hour, and incubated with specific antibodies in Can Get Signal immunoreaction enhancer solution (TOYOBO) at 4°C overnight. Endogenous Dicer and LGP2 were detected by specific antibodies against each protein (anti-Dicer, anti-LGP2 antibody), which were generated by immunizing rabbits with a synthetic peptide (*35*, *36*). TRBP and GFP were detected by anti-Myc antibody (Cell Signaling). The membranes were washed with TBS-T three times for 10 minutes each and reacted with HRP-linked anti-rabbit or anti-mouse antibody (GE Healthcare) at room temperature for 1 hour. The membranes were washed with TBS-T three times for 10 minutes each, incubated with ECL Prime Western Blotting Detection Reagent (GE Healthcare), and analyzed using the ImageQuant LAS4000 (GE Healthcare).

### Preparation of RNA-sequencing library

WT and TRBP^−/−^ HeLa cells were plated at 4 x 10^6^ cells / dish into a 9-cm dish and transfected with 6 µg/dish poly(I:C) using Lipofectamine 2000 (Invitrogen). At 0, 6, 8, and 10 h after transfection, cells were harvested and total RNAs were extracted. Extracted RNAs were measured by Qubit2.0 Fluorometer (Life Technologies) and the qualities were confirmed by Bioanalyzer (Agilent). rRNA was removed by Ribo-Zero rRNA Removal kit (AR BROWN). RNA-Seq was carried out using cDNA libraries prepared from rRNA-removed total RNA and small RNA fraction (15-110 nt) prepared by illumina Truseq small RNA kit. Total RNA is sequenced 36 nt by HiSeq 2500 in single-end mode and small RNA sequenced 100 nt by HiSeq 2500 in paired-end mode.

### RNA-sequencing analysis

For miRNA analysis, 16–28 nucleotide (nt)-long reads were extracted from small RNA sequence data as mature miRNAs using cutadapt (*37*)(version 1.11). The quality of all reads was confirmed to be sufficiently high using FastQC (http://www.bioinformatics.babraham.ac.uk/projects/fastqc, version 0.11.5). The 16–28 nt-long reads were mapped to RefSeq sequences (GRCh38) using Bowtie (*38*)(version 1.1.2), then annotated according to the miRNA database (*19–21*) (miRBase release 21). After counting raw reads using HTSeq (*39*)(version 0.7.2), reads corresponding to miRNAs were normalized by read per million (RPM). Expressed miRNAs were defined as miRNAs whose RPM at each time point was 1 or more. Upregulated miRNAs at each time point were defined as miRNAs whose RPM at each time point was 2-fold against 0h or more. Downregulated miRNAs at each time point were defined as miRNAs whose RPM at each time point was 0.5-fold or less against 0h. Relations between miRNAs and their host genes were identified by miRAID database (*24*)(https://www.miriad-database.org).

The target genes of each miRNA were predicted using TargetScan (*25*) (http://www.targetscan.org/vert_80/, release 8.0). The predicted targets were then defined by TargetScan cumulative context++ score (less than -0.25), which was calculated by summing up weighted context++ score obtained from conserved and nonconserved prediction files. Target genes of transcription factors were predicted using TFlink database (*28*) (https://tflink.net/) which estimates the transcriptional regulatory interactions from multiple experiments and databases. Gene ontology analysis was performed using DAVID web server (*26*, *27*) (https://david.ncifcrf.gov/home.jsp, v2024q1).

For mRNA analysis, low-quality bases (quality value < 30) were trimmed at the 3’ end of reads, and reads longer than 20 nt were extracted using cutadapt. An index for mapping was built from RefSeq sequences categorized as mature mRNAs with accession numbers prefixed by “NM_”, and reads were mapped using Bowtie. After counting raw reads using HTSeq, reads corresponding to transcripts were normalized by RPM. A gene expression network of miRNAs and their target genes was generated using Cytoscape(*40*) (version 3.10.1).

### Data Availability

RNA sequence data will be accessible via DDBJ Sequence Read Archive of National Institute of Genetics in Japan.

## Acknowledgment

S.A., Y.N., T.T., Y.A. and K.U.-T. designed the study at first, and S.A., Y.N., T.T., K.O, M.Y., Y.A. and K.U.-T. discussed the experimental procedures and results. S.A. and Y.N. performed the experiments and analyses of RNA sequence data. Y.S. performed RNA sequence, T.H. and H.I. designed miRNA inhibitors, Tough Decoys. The manuscript was drafted by S.A., Y.N., Y.A. and K.U.-T., and K.U.-T. reviewed the manuscript. All authors read and approved the final manuscript.

---

**Fig. S1:**
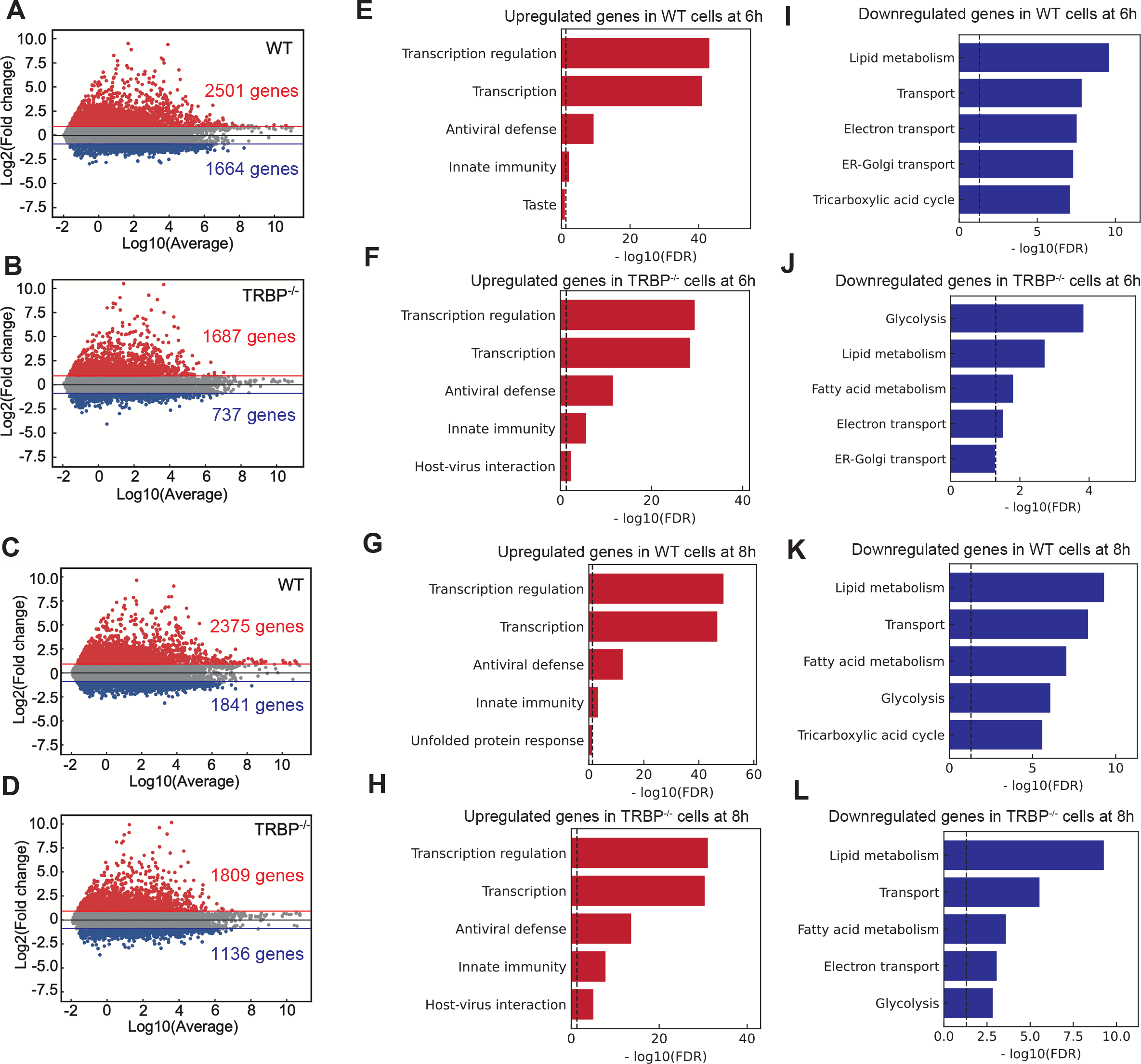
GO analysis of upregulated or downregulated genes by poly(I:C) transfection. (A, B) MA plot which shows changes of gene expression profile at 6 hours after poly(I:C) transfection in WT (A) or TRBP-/- (B) cells, respectively. Total of 15016 genes with RPKM > 0.1 were plotted. **(C, D)** MA plot which shows changes of gene expression profile at 8 hours after poly(I:C) transfection in WT (C) or TRBP-/- (D) cells, respectively. Total of 15016 genes with RPKM > 0.1 were plotted. **(E, F)** Top 5 GO terms, enriched in upregulated genes at 6 hours after poly(I:C) transfection in WT (E) or TRBP-/- (F) cells, respectively. **(G, H)** Top 5 GO terms which were enriched in upregulated genes at 8 hours after poly(I:C) transfection in WT (G) or TRBP-/- (H) cells, respectively. **(I, J)** Top 5 GO terms which were enriched in downregulated genes at 6 hours after poly(I:C) transfection in WT (I) or TRBP-/- (J) cells, respectively. **(K, L)** Top 5 GO terms which were enriched in downregulated genes at 8 hours after poly(I:C) transfection in WT (K) or TRBP-/- (L) cells, respectively.

**Fig. S2:**
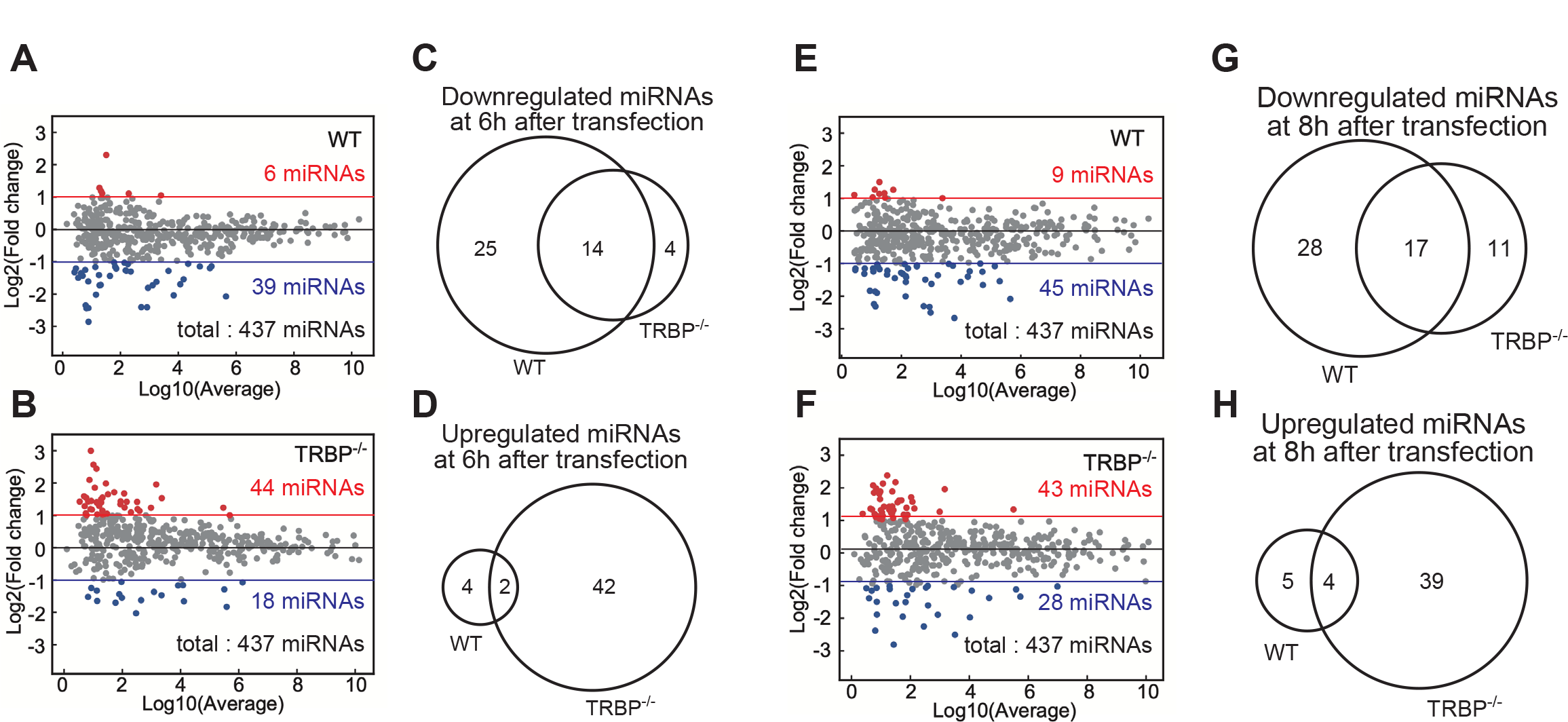
Changes in expression profiles of mature miRNAs by poly(I:C) transfection. (A, B) MA plot which shows changes of mature miRNA expression levels at 6 hours after poly(I:C) transfection in WT (A) or TRBP-/- (B) cells, respectively. **(C, D)** Venn’ s diagrams which show overlaps between downregulated (C) or upregulated (D) miRNAs in WT and TRBP-/- cells at 6 hours after poly(I:C) transfection. **(E, F)** MA plot which shows changes of mature miRNA expression levels at 8 hours after poly(I:C) transfection in WT (E) or TRBP-/- (F) cells, respectively. **(G, H)** Venn’ s diagrams which show overlaps between downregulated (G) or upregulated (H) miRNAs in WT and TRBP-/- cells at 8 hours after poly(I:C) transfection.

**Fig. S3:**
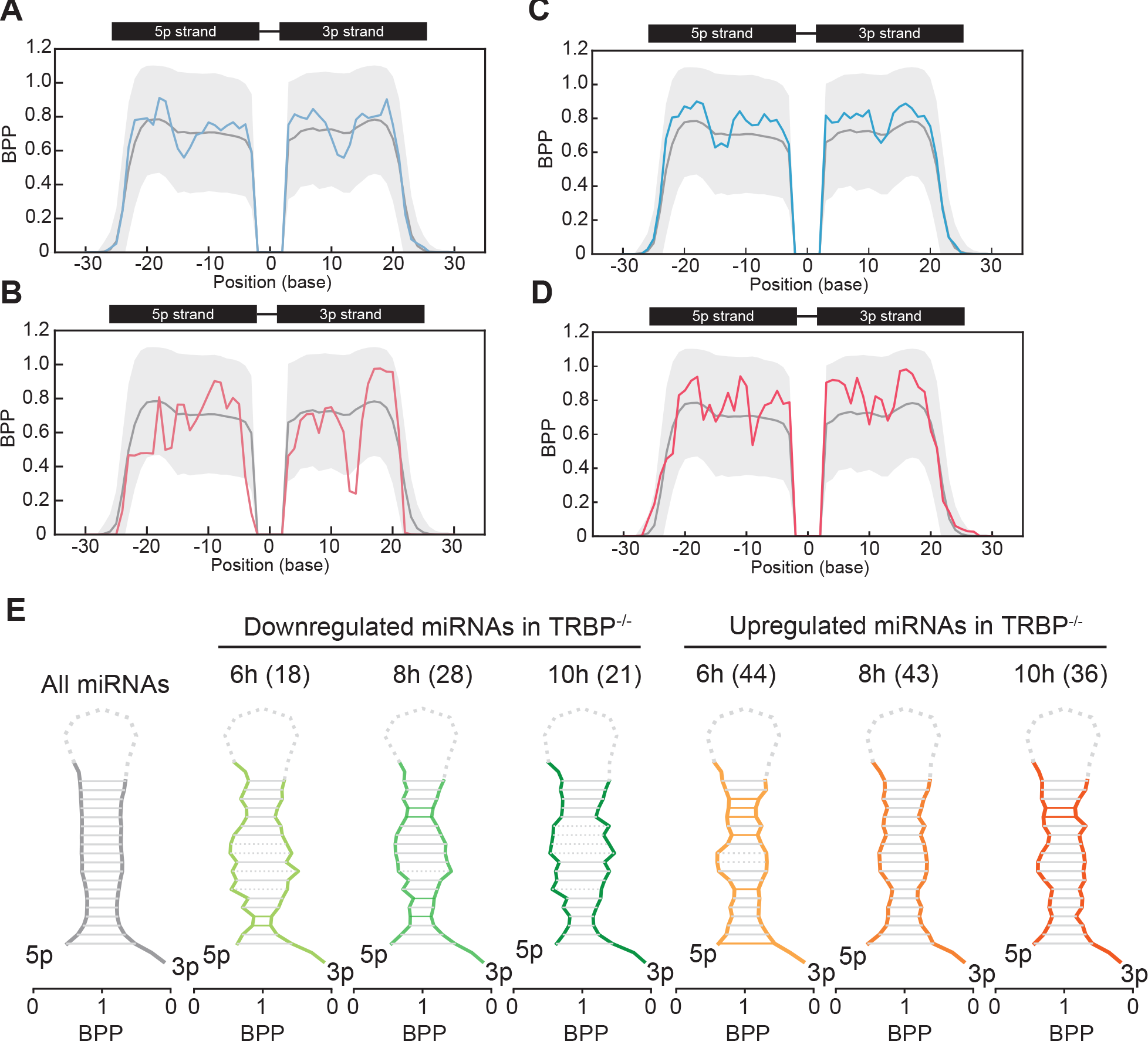
Base-pairing probabilities in precursor transcripts of mature miRNAs with altered expression level. (A-D) BPP values in the registered pre-miRNA structures of downregulated (A, C) or upregulated (B, D) mature miRNAs were calculated using the max value of base-pairing probabilities calculated by CentroidFold. The gray line indicates the mean BPP at each position of all pre-miRNAs registered in miRBase, and the gray area indicates the standard deviation. The average BPP of miRNAs which was downregulated only in WT cells was shown in blue (A) and those upregulated only in WT cells were shown in red (B) at 6 hours after poly(I:C) transfection. Also, BPP of miRNAs which was downregulated only in WT cells was shown in blue (C) and those upregulated only in WT cells were shown in red (D) at 8 hours after poly(I:C) transfection. **(E)** Predicted secondary structure of all pre-miRNAs (gray), downregulated miRNAs (green) and upregulated miRNAs (orange) in TRBP-/- cells reconstructed using BPP values. The stem-loop structure of pre-miRNA was created from a 5p strand and a 3p strand sequences with two nucleotide overhangs. Solid lines of blue or red show the positions with significantly higher BPPs determined by Student’ s t-test (*P < 0.05).

**Fig. S4:**
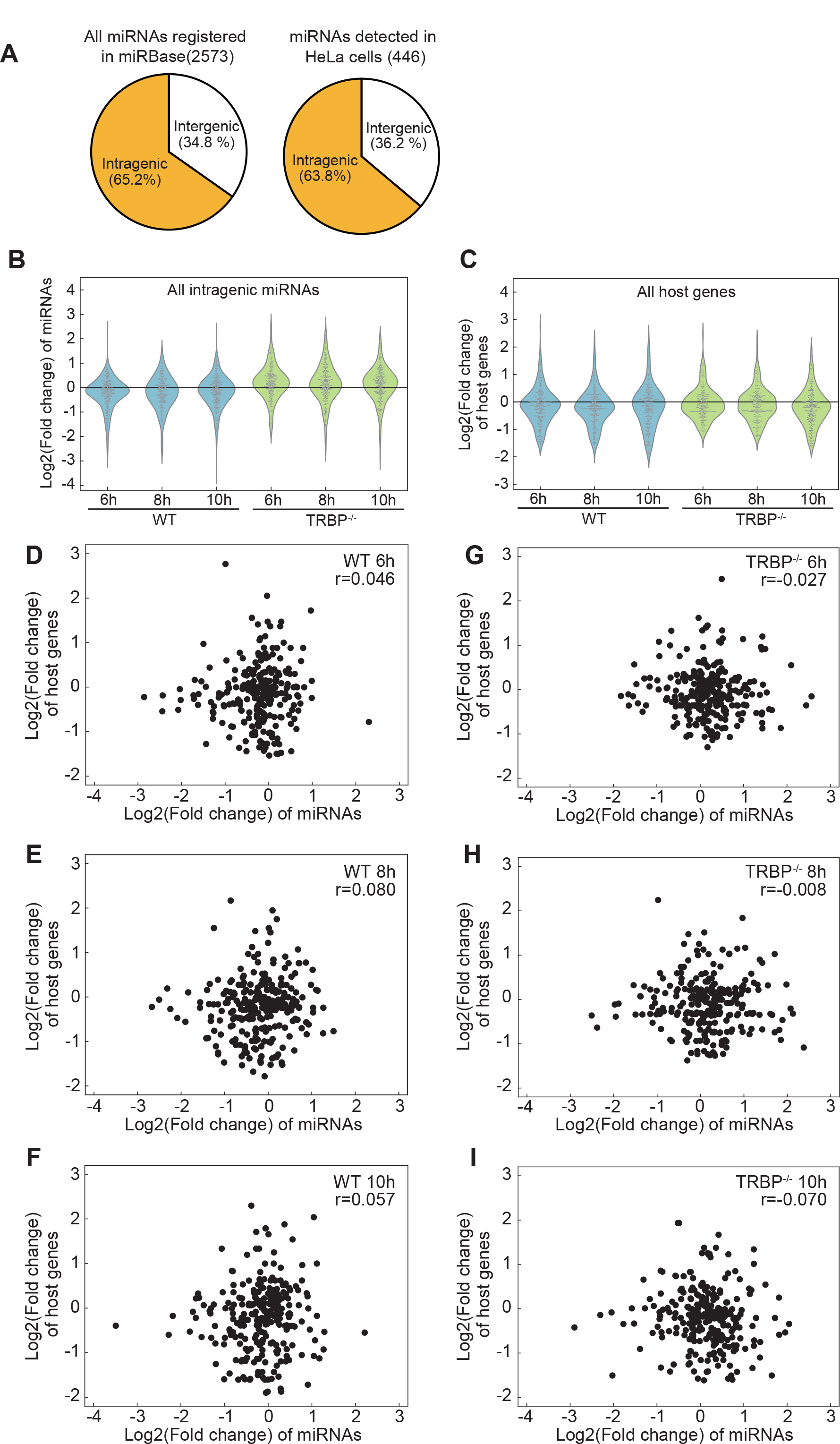
Correlation of expression levels between mature miRNAs and their host genes. (A) Percentage of intragenic (coded in exon or intron of a gene) miRNAs in all miRNAs registered in miRbase (left) and expressed (RPM > 1 in all sample) miRNAs in small RNA-seq (right). **(B)** Violin plots which show log2(fold change) distribution of intragenic miRNAs in WT and TRBP-/- cells at each time point. **(C)** Violin plots which show log2(fold change) distribution of host genes of miRNAs in WT and TRBP-/- cells at each time point. **(D-I)** XY plots which show correlations between the intragenic mature miRNA levels and the expression levels of their host genes. X axis indicates log2(fold change)s of miRNAs. Y axis, those of host genes.

**Fig. S5:**
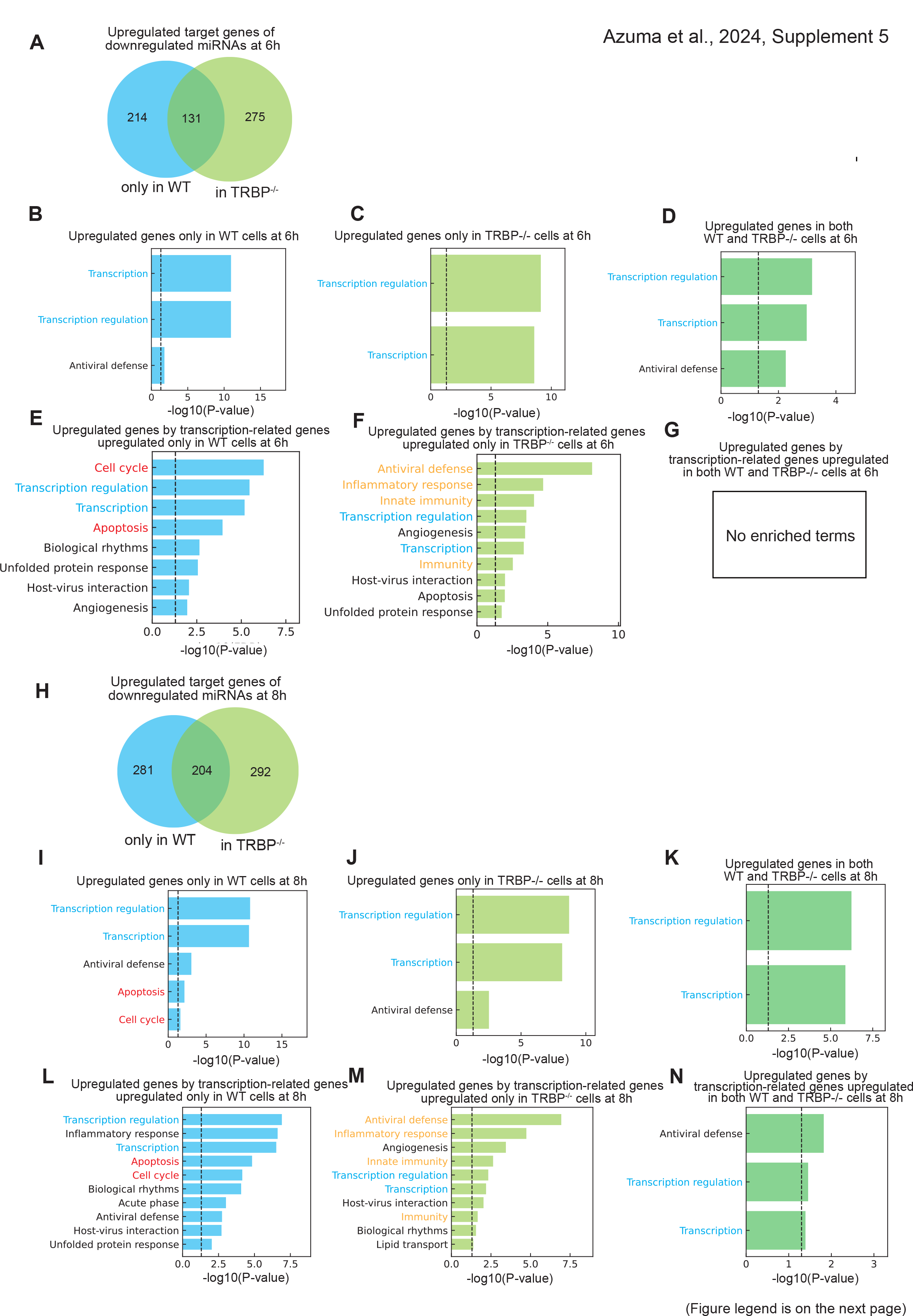
Predicted target genes regulated by downregulated mature miRNAs or upregulated transcription factors during poly(I:C) transfection. (A) Venn’ s diagram indicating the overlap between upregulated genes by downregulated miRNAs in only WT cells and in TRBP-/- cells at 6 hours after poly(I:C) transfection. **(B)** Enriched GO terms in upregulated genes only in WT cells at 6 hours after poly(I:C) transfection (sky-blue region in Fig. S5A). **(C)** Enriched GO terms in upregulated genes only in TRBP-/- cells at 6 hours after poly(I:C) transfection (light-green region in Fig. S5A). **(D)** Enriched GO terms in upregulated genes both in WT and TRBP-/- cells at 6 hours after poly(I:C) transfection (green region in Fig. S5A). **(E)** Enriched GO terms in upregulated target genes by transcription-related genes, specifically upregulated only in WT cells at 6 hours (Term “Transcription” in Fig. S5B). **(F)** Enriched GO terms in upregulated target genes by transcription factors, specifically upregulated only in TRBP-/- cells at 6 hours (Term “Transcription” in Fig. S5C). **(G)** Enriched GO terms in upregulated target genes by transcription factors, commonly upregulated both in WT and TRBP-/- cells at 6 hours (Term “Transcription” in Fig. S5D). **(H)** Venn’ s diagram indicating the overlap between upregulated genes by downregulated miRNAs in only WT cells and in TRBP-/- cells at 8 hours after poly(I:C) transfection. **(I)** Enriched GO terms in upregulated genes only in WT cells at 8 hours after poly(I:C) transfection (sky-blue region in Fig. S5H). **(J)** Enriched GO terms in upregulated genes only in TRBP-/- cells at 8 hours after poly(I:C) transfection (light-green region in Fig. S5H). **(K)** Enriched GO terms in upregulated genes both in WT and TRBP-/- cells at 8 hours after poly(I:C) transfection (green region in Fig. S5H). **(L)** Enriched GO terms in upregulated target genes by transcription-related genes, specifically upregulated only in WT cells at 8 hours (Term “Transcription” in Fig. S5I). **(M)** Enriched GO terms in upregulated target genes by transcription factors, specifically upregulated only in TRBP-/- cells at 8 hours (Term “Transcription” in Fig. S5J). **(N)** Enriched GO terms in upregulated target genes by transcription factors, commonly upregulated both in WT and TRBP-/- cells at 8 hours (Term “Transcription” in Fig. S5K).

**Fig. S6:**
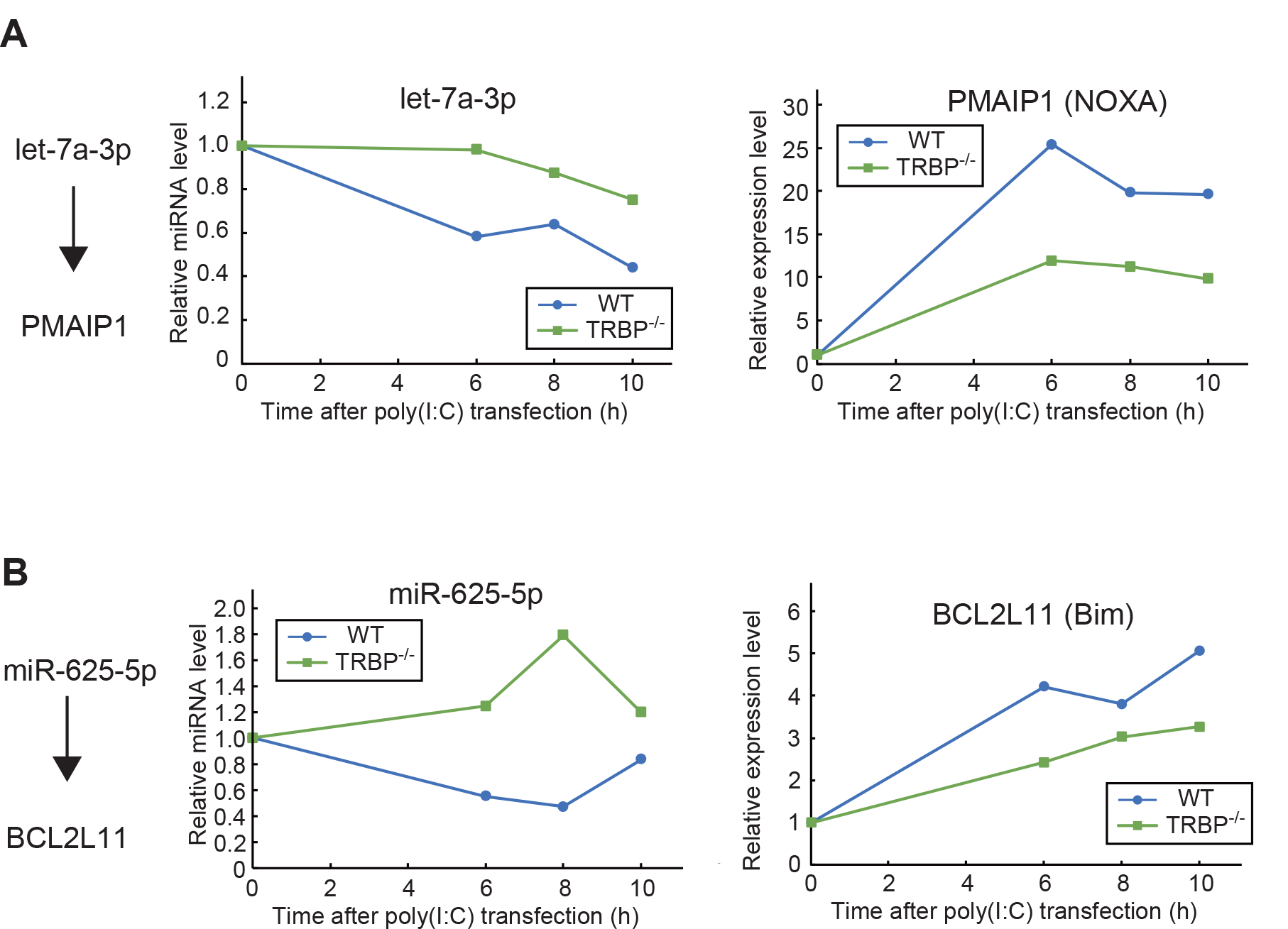
Expression levels of mature miRNAs and apoptosis-related genes. (A) Relative miRNA levels of let-7a-5p (left) and mRNA level of PMAIP1 (right) which was a predicted target of let-7a-3p. RPMs or RPKMs at 6, 8 10 hours after poly(I:C) transfection were divided by that of 0 hours in WT cells (blue) and TRBP-/- cells (green) respectively. **(B)** Relative miRNA levels of miR-625-5p (left) and mRNA level of BCL2L11 (right) which was a predicted target of miR-625-5p. RPMs or RPKMs at 6, 8 10 hours after poly(I:C) transfection were divided by that of 0 hours in WT cells (blue) and TRBP-/- cells (green) respectively.

**Table S1.**
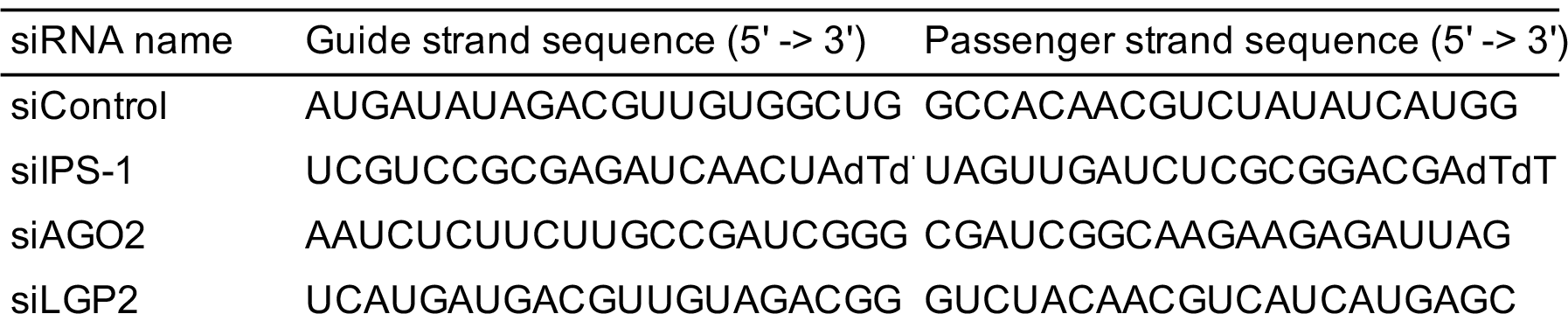
siRNA sequences used for gene knockdown.

**Table S2.**
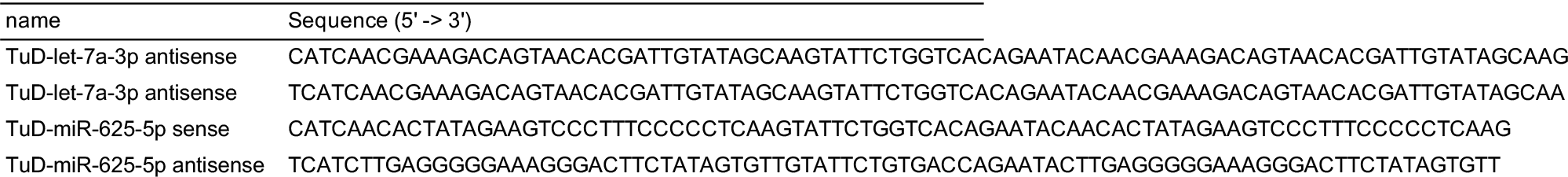
Oligonucleotides used for the construction of plasmids expressing miRNA inhibitors (TuD-decoys).

**Table S3.**
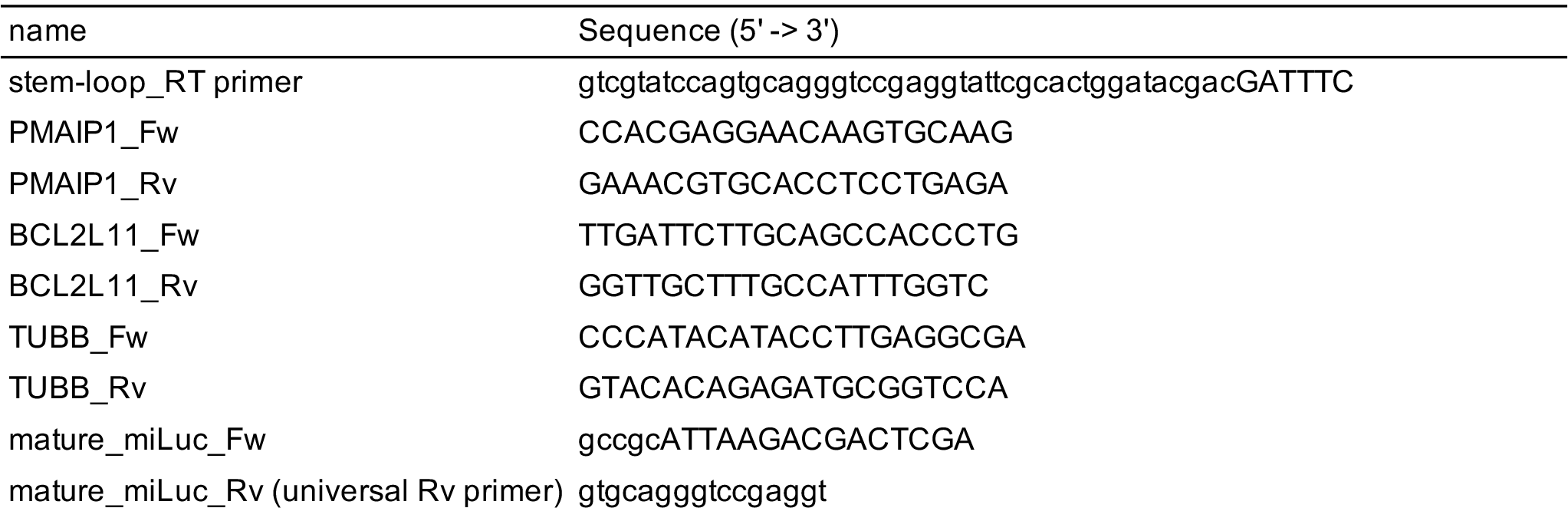
Oligonucleotides used for qRT/RT-PCR.

## References

1. Y. Lee, M. Kim, J. Han, K.-H. Yeom, S. Lee, S. H. Baek, V. N. Kim, MicroRNA genes are transcribed by RNA polymerase II. EMBO J. 23, 4051–4060 (2004).

2. Y. Lee, C. Ahn, J. Han, H. Choi, J. Kim, J. Yim, J. Lee, P. Provost, O. Rådmark, S. Kim, V. N. Kim, The nuclear RNase III Drosha initiates microRNA processing. Nature 425, 415– 419 (2003).

3. R. Yi, Y. Qin, I. G. Macara, B. R. Cullen, Exportin-5 mediates the nuclear export of pre- microRNAs and short hairpin RNAs. Genes Dev. 17, 3011–3016 (2003).

4. M. T. Bohnsack, K. Czaplinski, D. Gorlich, Exportin 5 is a RanGTP-dependent dsRNA- binding protein that mediates nuclear export of pre-miRNAs. RNA 10, 185–191 (2004).

5. P. Provost, D. Dishart, J. Doucet, D. Frendewey, B. Samuelsson, O. Rådmark, Ribonuclease activity and RNA binding of recombinant human Dicer. EMBO J. 21, 5864–5874 (2002).

6. T. P. Chendrimada, R. I. Gregory, E. Kumaraswamy, J. Norman, N. Cooch, K. Nishikura, R. Shiekhattar, TRBP recruits the Dicer complex to Ago2 for microRNA processing and gene silencing. Nature 436, 740–744 (2005).

7. I. J. MacRae, E. Ma, M. Zhou, C. V. Robinson, J. A. Doudna, In vitro reconstitution of the human RISC-loading complex. Proc. Natl. Acad. Sci. U. S. A. 105, 512–517 (2008).

8. E. Maniataki, Z. Mourelatos, A human, ATP-independent, RISC assembly machine fueled by pre-miRNA. Genes Dev. 19, 2979–2990 (2005).

9. H.-W. Wang, C. Noland, B. Siridechadilok, D. W. Taylor, E. Ma, K. Felderer, J. A. Doudna, E. Nogales, Structural insights into RNA processing by the human RISC-loading complex. Nat. Struct. Mol. Biol. 16, 1148–1153 (2009).

10. T. Takahashi, Y. Nakano, K. Onomoto, M. Yoneyama, K. Ui-Tei, LGP2 virus sensor enhances apoptosis by upregulating apoptosis regulatory genes through TRBP-bound miRNAs during viral infection. Nucleic Acids Res. 48, 1494–1507 (2020).

11. L. B. Ivashkiv, L. T. Donlin, Regulation of type I interferon responses. Nat. Rev. Immunol. 14, 36–49 (2014).

12. M. Yoneyama, M. Kikuchi, T. Natsukawa, N. Shinobu, T. Imaizumi, M. Miyagishi, K. Taira, S. Akira, T. Fujita, The RNA helicase RIG-I has an essential function in double-stranded RNA-induced innate antiviral responses. Nat. Immunol. 5, 730–737 (2004).

13. M. Yoneyama, M. Kikuchi, K. Matsumoto, T. Imaizumi, M. Miyagishi, K. Taira, E. Foy, Y.-M. Loo, M. Gale Jr, S. Akira, S. Yonehara, A. Kato, T. Fujita, Shared and unique functions of the DExD/H-box helicases RIG-I, MDA5, and LGP2 in antiviral innate immunity. J. Immunol. 175, 2851–2858 (2005).

14. T. Takahashi, Y. Nakano, K. Onomoto, F. Murakami, C. Komori, Y. Suzuki, M. Yoneyama, K. Ui-Tei, LGP2 virus sensor regulates gene expression network mediated by TRBP-bound microRNAs. Nucleic Acids Res. 46, 9134–9147 (2018).

15. S. Chakravarthy, S. H. Sternberg, C. A. Kellenberger, J. A. Doudna, Substrate-Specific Kinetics of Dicer-Catalyzed RNA Processing. J. Mol. Biol. 404, 392–402 (2010).

16. A. Daher, M. Longuet, D. Dorin, F. Bois, E. Segeral, S. Bannwarth, P. L. Battisti, D. F. Purcell, R. Benarous, C. Vaquero, E. F. Meurs, A. Gatignol, Two dimerization domains in the trans-activation response RNA-binding protein (TRBP) individually reverse the protein kinase R inhibition of HIV-1 long terminal repeat expression. J. Biol. Chem. 276, 33899– 33905 (2001).

17. A. García-Sastre, Ten Strategies of Interferon Evasion by Viruses. Cell Host Microbe 22, 176–184 (2017).

18. S. H. Kaufmann, S. Desnoyers, Y. Ottaviano, N. E. Davidson, G. G. Poirier, Specific proteolytic cleavage of poly(ADP-ribose) polymerase: an early marker of chemotherapy- induced apoptosis. Cancer Res. 53, 3976–3985 (1993).

19. S. Griffiths-Jones, The microRNA Registry. Nucleic Acids Res. 32, D109–11 (2004).

20. S. Griffiths-Jones, R. J. Grocock, S. van Dongen, A. Bateman, A. J. Enright, miRBase: microRNA sequences, targets and gene nomenclature. Nucleic Acids Res. 34, D140–D144 (2006).

21. A. Kozomara, S. Griffiths-Jones, miRBase: annotating high confidence microRNAs using deep sequencing data. Nucleic Acids Res. 42, D68–73 (2014).

22. S. L. Ameres, M. D. Horwich, J.-H. Hung, J. Xu, M. Ghildiyal, Z. Weng, P. D. Zamore, Target RNA-directed trimming and tailing of small silencing RNAs. Science 328, 1534– 1539 (2010).

23. B. Liu, Y. Shyr, J. Cai, Q. Liu, Interplay between miRNAs and host genes and their role in cancer. Brief. Funct. Genomics 18, 255–266 (2019).

24. L. C. Hinske, G. S. França, H. A. M. Torres, D. T. Ohara, C. M. Lopes-Ramos, J. Heyn, L. F. L. Reis, L. Ohno-Machado, S. Kreth, P. A. F. Galante, miRIAD-integrating microRNA inter- and intragenic data. Database 2014 (2014).

25. V. Agarwal, G. W. Bell, J.-W. Nam, D. P. Bartel, Predicting effective microRNA target sites in mammalian mRNAs. Elife 4 (2015).

26. D. W. Huang, B. T. Sherman, R. A. Lempicki, Systematic and integrative analysis of large gene lists using DAVID bioinformatics resources. Nat. Protoc. 4, 44–57 (2009).

27. B. T. Sherman, M. Hao, J. Qiu, X. Jiao, M. W. Baseler, H. C. Lane, T. Imamichi, W. Chang, DAVID: a web server for functional enrichment analysis and functional annotation of gene lists (2021 update). Nucleic Acids Res. 50, W216–W221 (2022).

28. O. Liska, B. Bohár, A. Hidas, T. Korcsmáros, B. Papp, D. Fazekas, E. Ari, TFLink: an integrated gateway to access transcription factor-target gene interactions for multiple species. Database 2022 (2022).

29. T. Haraguchi, Y. Ozaki, H. Iba, Vectors expressing efficient RNA decoys achieve the long- term suppression of specific microRNA activity in mammalian cells. Nucleic Acids Res. 37, e43 (2009).

30. S. Ma, A. Kotar, I. Hall, S. Grote, S. Rouskin, S. C. Keane, Structure of pre-miR-31 reveals an active role in Dicer–TRBP complex processing. Proceedings of the National Academy of Sciences 120, e2300527120 (2023).

31. T. Takahashi, S. Zenno, O. Ishibashi, T. Takizawa, K. Saigo, K. Ui-Tei, Interactions between the non-seed region of siRNA and RNA-binding RLC/RISC proteins, Ago and TRBP, in mammalian cells. Nucleic Acids Res. 42, 5256–5269 (2014).

32. Y. Kim, J. Yeo, J. H. Lee, J. Cho, D. Seo, J.-S. Kim, V. N. Kim, Deletion of Human tarbp2 Reveals Cellular MicroRNA Targets and Cell-Cycle Function of TRBP. Cell Rep. 9, 1061– 1074 (2014).

33. J. Chen, The Cell-Cycle Arrest and Apoptotic Functions of p53 in Tumor Initiation and Progression. Cold Spring Harb. Perspect. Med. 6, a026104 (2016).

34. T. Takahashi, T. Miyakawa, S. Zenno, K. Nishi, M. Tanokura, K. Ui-Tei, Distinguishable in vitro binding mode of monomeric TRBP and dimeric PACT with siRNA. PLoS One 8, e63434 (2013).

35. N. Doi, S. Zenno, R. Ueda, H. Ohki-Hamazaki, K. Ui-Tei, K. Saigo, Short-interfering-RNA- mediated gene silencing in mammalian cells requires Dicer and eIF2C translation initiation factors. Curr. Biol. 13, 41–46 (2003).

36. K. Onomoto, M. Jogi, J.-S. Yoo, R. Narita, S. Morimoto, A. Takemura, S. Sambhara, A. Kawaguchi, S. Osari, K. Nagata, T. Matsumiya, H. Namiki, M. Yoneyama, T. Fujita, Critical role of an antiviral stress granule containing RIG-I and PKR in viral detection and innate immunity. PLoS One 7, e43031 (2012).

37. M. Martin, Cutadapt removes adapter sequences from high-throughput sequencing reads. EMBnet.journal 17, 10–12 (2011).

38. B. Langmead, C. Trapnell, M. Pop, S. L. Salzberg, Ultrafast and memory-efficient alignment of short DNA sequences to the human genome. Genome Biol. 10, R25 (2009).

39. S. Anders, P. T. Pyl, W. Huber, HTSeq--a Python framework to work with high-throughput sequencing data. Bioinformatics 31, 166–169 (2015).

40. P. Shannon, A. Markiel, O. Ozier, N. S. Baliga, J. T. Wang, D. Ramage, N. Amin, B. Schwikowski, T. Ideker, Cytoscape: a software environment for integrated models of biomolecular interaction networks. Genome Res. 13, 2498–2504 (2003).

